# Gene inversion triggered origination of brackish archaeal heterotrophs in the aftermath of the Cryogenian Snowball Earth

**DOI:** 10.1101/2022.09.25.509439

**Authors:** Lu Fan, Bu Xu, Songze Chen, Yang Liu, Fuyan Li, Wei Xie, Apoorva Prabhu, Dayu Zou, Ru Wan, Hongliang Li, Haodong Liu, Yuhang Liu, Shuh-Ji Kao, Jianfang Chen, Yuanqing Zhu, Christian Rinke, Meng Li, Maoyan Zhu, Chuanlun Zhang

## Abstract

Land-ocean interactions greatly impacted the evolution of coastal life on Earth. However, the geological forces and genetic mechanisms that shaped evolutionary adaptations and allowed microorganisms to inhabit coastal brackish waters remain largely unexplored. Here, we infer the evolutionary trajectory of the ubiquitous heterotrophic archaea *Poseidoniales* (Marine Group II archaea) across global aquatic habitats. Our results show that their brackish subgroups have evolved through the rearrangement of the magnesium transport gene *corA* that conferred osmotic-stress tolerance dated to over 600 million years ago. The coastal family of *Poseidoniales* showed a rapid increase in the evolutionary rate during and in the aftermath of the Cryogenian Snowball Earth (~700 million years ago), possibly in response to the enhanced phosphorus supply and the rise of algae. Our study highlights the close interplay between genetic changes and ecosystem transformations that boosted microbial diversification in the Neoproterozoic continental margins.

## Introduction

Salinity is among the strongest environmental factors determining community composition of both macro- and micro-organisms (Lee and Bell, 1999; Lozupone and Knight, 2007), which is so called the ‘salinity divide’ (Logares et al., 2009). Compared to distinct and characteristic biological populations in marine and freshwater ecosystems, biodiversity in brackish environments such as estuaries and enclosed seas is less understood despite the importance of these environments in land-ocean interactions and their vulnerability to human activities. Recently, the existence of a unique, highly diverse and possibly globally distributed brackish community of bacteria and protists has been reported (Filker et al., 2019; Herlemann et al., 2011; Hugerth et al., 2015). Two very recent studies inferred that the evolutionary transitions of bacteria and archaea between freshwater, brackish and marine environments started in the early Earth (Jurdzinski et al., 2023; Ngugi et al., 2023); yet the origination of brackish microorganisms is still largely unknown.

Brackish ecosystems are different from marine and freshwater ones in various environmental factors such as salinity, nutrients, predators and competitors (Walsh et al., 2013). It is thus challenging to identify the key genetic changes and the primary selective force triggering the divergence between these ecosystems. Recent genome comparisons among saline and freshwater subgroups of bacteria and archaea have revealed functional differences in these two subgroups in osmotic regulation and in growth substrate specificity (Cabello-Yeves et al., 2017; Eiler et al., 2016; Martijn et al., 2020; Penn and Jensen, 2012; Ren and Wang, 2022; Simon et al., 2017; H. Zhang et al., 2019). Two fundamental questions are 1) the primary versus the secondary genetic changes during the transition and 2) whether the transition was caused by a sudden event or by gradual and cumulative effects (Eiler et al., 2016; Henson et al., 2018).

Land-ocean interactions may have facilitated marine life evolution in geological history of the earth. A typical example is the Cambrian explosion of animals, which happened largely on continental margins (Wood et al., 2019). This drastic shift of global biota is attributed to interconnected global changes after the Cryogenian Snowball Earth and in the Ediacaran-Cambrian transition (Smith and Harper, 2013; Wood et al., 2019) (~700 – 485.4 million years ago, Ma), with rapid sea level rise, orogeny and enhanced land weathering (Halverson et al., 2010; Sharoni and Halevy, 2023). Consequently, pulses of nutrients supply to the ocean (Reinhard et al., 2017) boosted coastal primary productivity (Xiang et al., 2018) and food web complexity (Butterfield, 2009). At the same time, the rise and variability of ocean oxygen level (Chen et al., 2015; F. Zhang et al., 2019) might accompanied the emergence of algae (Brocks et al., 2017). While numerous fossil records provide details in the radiation of multicellular animals in this period (Erwin and Valentine, 2013), evidence of change in evolution and ecology of bacterial and archaeal heterotrophs is lacking. Specifically, it is unclear whether the microbial evolutionary rate accelerated as their animal counterparts did (Erwin et al., 2011), and whether changes in quantity and quality of organic matter altered the ‘microbial loop’ (Pehr et al., 2018) of the transforming food webs in the Ediacaran-Cambrian coastal waters. Moreover, the postglacial ocean is considered to be stably stratified with a surface layer of brackish and oxic water (Hurtgen et al., 2006) but the biological impact of these expanded brackish environments to microbial evolution is currently unknown.

Marine Group II archaea (*Ca*. *Poseidoniales*) are among the most abundant planktonic archaea in global oceans with great ecological potentials (Rinke et al., 2019; Tully, 2019). Genomic analyses support that they are heterotrophs living on algal-derived organic substrates and are an important part of the ‘microbial loop’ (Damashek et al., 2021; Orellana et al., 2019; Pereira et al., 2020; Rinke et al., 2019; Tully, 2019; Zhang et al., 2015) and the microbial carbon pump processes (Zhang et al., 2018) in the open ocean and coastal marine environments. High abundance of *Poseidoniales* 16S rRNA genes was recently reported at the estuary mixing zone with salinity below 15‰ (Xie et al., 2017), suggesting the possible existence of brackish *Poseidoniales* populations. In this study, an in-depth analysis of global brackish metagenomes allowed us to identify the evolutionary transition of *Poseidoniales* between marine and brackish habitats and to elucidate the important genetic changes in the origination of the brackish subclades. Furthermore, the two families of *Poseidoniales*, which adapted to nearshore and pelagic waters, respectively, were carefully compared in evolutionary rate to reveal the possible geological and environmental driving forces during this marine-brackish shift.

## Results and Discussion

### Global distribution of diverse and active brackish-specific *Poseidoniaceae*

Metagenome-assembled genomes (MAGs) of *Poseidoniales* were reconstructed from 282 metagenomes of global estuarine, enclosed sea and coastal environments (fig. S1 and table S1) and phylogenetically analyzed together with the ones obtained in recent studies (Orellana et al., 2019; Rinke et al., 2019; Tully, 2019). This approach contributed 94 (20.7%) high-quality novel genomes to the updated non-redundant *Poseidoniales* genome dataset (455 MAGs, completeness median = 84.36%, contamination median = 1.09%) based on a cutoff of 99% average nucleotide identity (ANI) (table S2), which would contribute to a better understanding of species diversity of *Poseidoniales* in low-salinity environments. *Poseidoniales* include two family-level subgroups, MGIIa (*Poseidoniaceae*) and MGIIb (*Thalassarchaeaceae*) (Rinke et al., 2019). Abundance mapping of this non-redundant MAG dataset in global surface waters with distinct salinities shows that *Poseidoniaceae* are in high abundance in enclosed seas and estuaries, while *Thalassarchaeaceae* are only detected in most of the coastal and open ocean samples (Fig. 1A). This spatial distribution pattern is in line with the previous observation that *Poseidoniaceae* are adapted to more eutrophic and diverse coastal environments but *Thalassarchaeaceae* usually stay in the open ocean (Damashek et al., 2021; Orellana et al., 2019; Pereira et al., 2020; Rinke et al., 2019; Tully, 2019; Zhang et al., 2015). Notably, two patterns of abundance distribution along the salinity gradient are detected in *Poseidoniaceae*: a ‘salt-preferred’ pattern in which their abundance increased with the increase of salinity and a ‘brackish specific’ pattern in which they were enriched in salinity between 6.6‰ and 23‰ but depleted or absent in salinity beyond this range (Fig. 1A). The metatranscriptomic analysis supports that *Poseidoniaceae* of both patterns were in an active state (fig. S2).

**Fig. 1.**
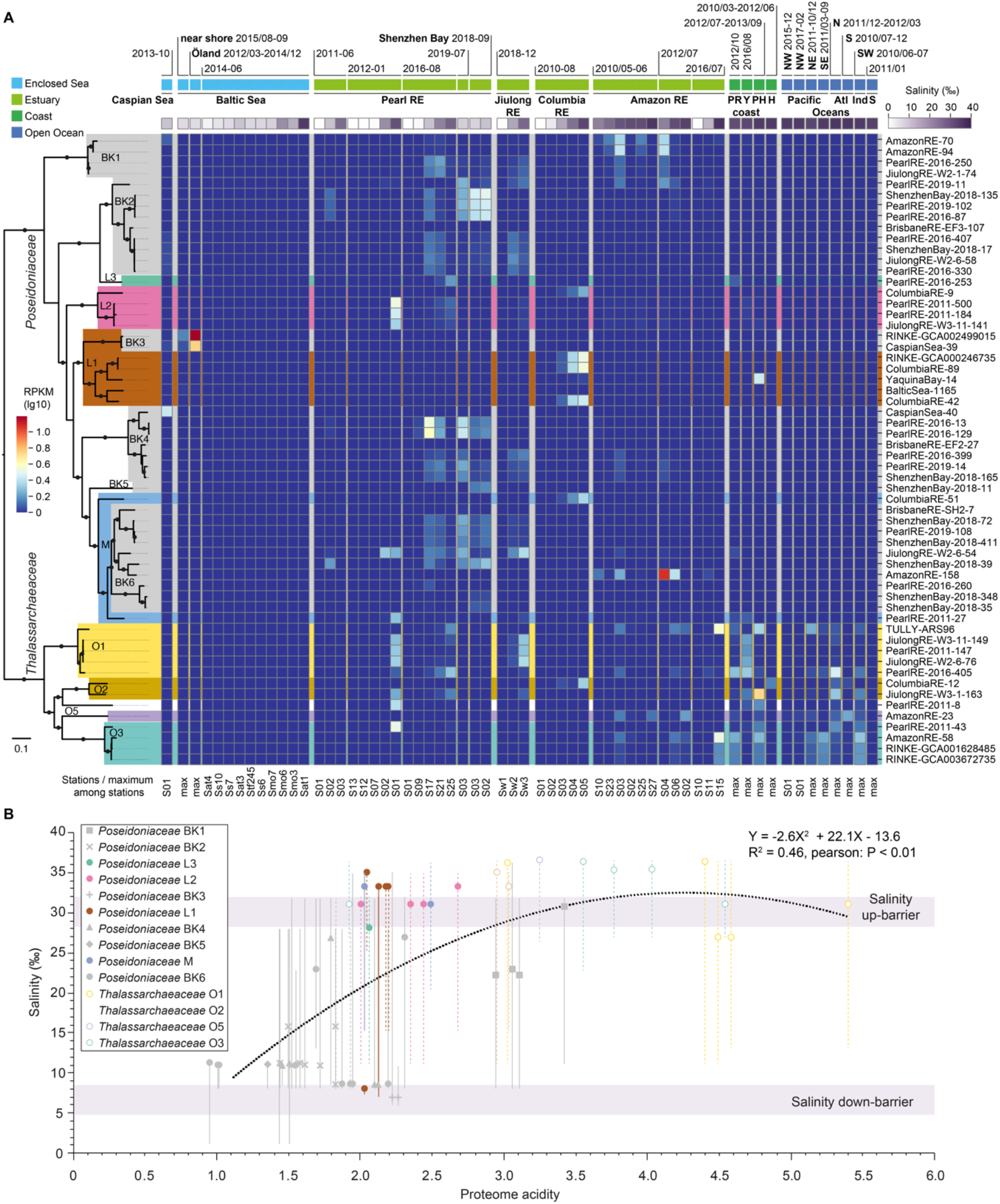
Global distribution and proteome adaptation of brackish-specific and marine-specific *Poseidoniales*. (**A**) Abundance pattern of *Poseidoniales* MAGs based on 246 metagenome samples from surface waters of global marine and brackish environments. MAGs with RPKM value above 0.4 in enclosed sea and estuarine samples are shown. In the maximum-likelihood tree at the left of the panel, solid dots on internal branches show branch supports of ultra-fast bootstrapping (1000) in IQTree > 90%. Shades on branches show *Poseidoniales* genera with the color code according to Rinke *et al*. 2019 (Rinke et al., 2019), except for brackish clades that are in grey. Abbreviations of sampling areas are: RE = river estuary, PR = Pearl River estuary, Y = Yangtze River estuary, PH = Port Hacking, H= Helgoland, Alt = Atlantic, Ind = Indian, S= Southern/South, N = North, NW = Northwest, NE = Northeast, SW = Southwest, SE= Southeast. Salinity shows the sample salinity or the average of the collection of samples. RPKM shows the abundance or the maximum abundance of each MAG in each sample or a collection of samples, respectively. (**B**) Habitat salinity and proteome acidity of *Poseidoniales* MAGs found in estuaries and enclosed seas. Circles show *Poseidoniales* MAGs. Filled circles and solid lines belong to *Poseidoniaceae* MAGs, while empty circles and dashed lines belong to *Thalassarchaeaceae* MAGs. The position of each dot on the y-axis shows the optimal salinity of that MAG. Regression of these dots is shown as the dashed black line. The scale of each vertical line on the y-axis shows the upper and down limits salinity of that MAG. The color code of dots and lines shows *Poseidoniales* genera according to Rinke *et al*. 2019 (Rinke et al., 2019), except for brackish clades which are in grey.

In the phylogenomic tree, *Poseidoniaceae* genomes from global estuaries and enclosed seas form several clusters (fig. S3). Based on the above abundance analysis, six monophyletic brackish-specific clades can be identified (Fig. 1A and fig. S3). They either branch within or as close sisters to previously classified *Poseidoniaceae* genera (Rinke et al., 2019). Some of these brackish-specific lineages are found in different estuaries globally, while others are found to be enriched only in one estuary or one enclosed sea (Fig. 1A), implying that global dispersion and local adaptation may both play a role in their geographic distribution (Jurdzinski et al., 2023). Repeated presence but interannual variability in abundance of these brackish specific lineages are observed in the Baltic Sea, the Pearl River estuary and the Amazon River estuary, suggesting they have specifically adapted to coastal brackish waters but factors other than salinity might impact their temporary abundances (Orellana et al., 2019). Notably, estuaries harbor a much greater diversity of brackish MAGs than enclosed seas demonstrating the importance of estuaries (possibly because of its environmental dynamics) in studying brackish microorganisms (Fig. 1A) (Jurdzinski et al., 2023).

Acidified proteome isoelectric point (p*I*) is recognized to be a strong indicator of microorganisms inhabiting saline environments (Cabello-Yeves and Rodriguez-Valera, 2019) as a response to higher intracellular ion concentration (Fagerbakke et al., 1999). To verify that the brackish-enriched *Poseidoniaceae* are specifically adapted to low-salinity habitats, we calculated the optimum salinity values of the MAGs (*i.e.*, the salinity of an environment in which a MAG has the highest abundance) detected in enclosed sea and estuarine samples and plotted them according to the estimated acidity of their proteome p*I* patterns. Fig. 1B clearly shows a polynomial correlation (R^2^ = 0.46, P < 0.01, n = 56) between proteome acidity and salinity adaptation. Almost all of *Thalassarchaeaceae* MAGs have acidity values over 2.9 (except one) and optimum salinity over 30‰ (except two). Most *Poseidoniaceae* with acidity above 2.0 enriched in salinity from 20‰ to 35‰, with an up-limit of over 36‰. In contrast, *Poseidoniaceae* MAGs of acidity below 2.0 belonging to brackish-specific clades have optimum salinities below 30‰ and down to 8‰, which are detectable even in river mouth at a salinity around 1‰ but never at a salinity above 32‰. This distinct distribution pattern of marine and brackish *Poseidoniaceae* subgroups is consistent with the suggestion of long-term divergent evolution (Cabello-Yeves and Rodriguez-Valera, 2019).

### *Poseidoniales* potentially caused by a *corA* gene inversion and establishment in a stress-response gene cluster

Researchers have proposed various genes potentially contributing to the land-ocean divergence in microbial evolution including those functioning in osmotic regulation, substrate preference and adaptation to dynamic environments (Jurdzinski et al., 2023; Logares et al., 2009). To identify genes potentially responsible for differentiating the marine and brackish *Poseidoniales* subgroups, we annotated *Poseidoniaceae* and *Thalassarchaeaceae* MAGs (table S2) and conducted gene-centered comparison. Remarkably, the magnesium transporter gene *corA* is the only one that is present in > 85% MAGs (30/33, 90.9%) of the brackish clades and in < 5% MAGs (18/369, 4.9%) of the marine clades of *Poseidoniales* (Fig. 2C and fig. S4). Student’s *t*-Test shows that the proteome acidities of *Poseidoniaceae* encoding *corA* are significantly lower (P = 0.016, two-tailed) than those without this gene (Fig. 2A). CorA is one of the main import channel of magnesium ions (Mg^2+^) in bacteria and archaea (Maguire, 2006). Its potential bidirectional and concentration-regulated feature (Payandeh et al., 2013) suggests it may act as a condition-dependent vale to maintain a stable intracellular magnesium concentration in hydro-dynamic estuaries facing stresses of sudden increasing or decreasing salinity. This observation suggests that intracellular magnesium, which can potential stabilize or modulate structures of macromolecules such as DNAs, RNAs, and proteins, may be important in adaptation of brackish *Poseidoniaceae* (Supplementary Text). It also highlights that the stress of salinity fluctuation in brackish environments instead of a steady and moderate salinity is potentially the primary barrier for brackish water adaptation of *Poseidoniales*.

**Fig. 2.**
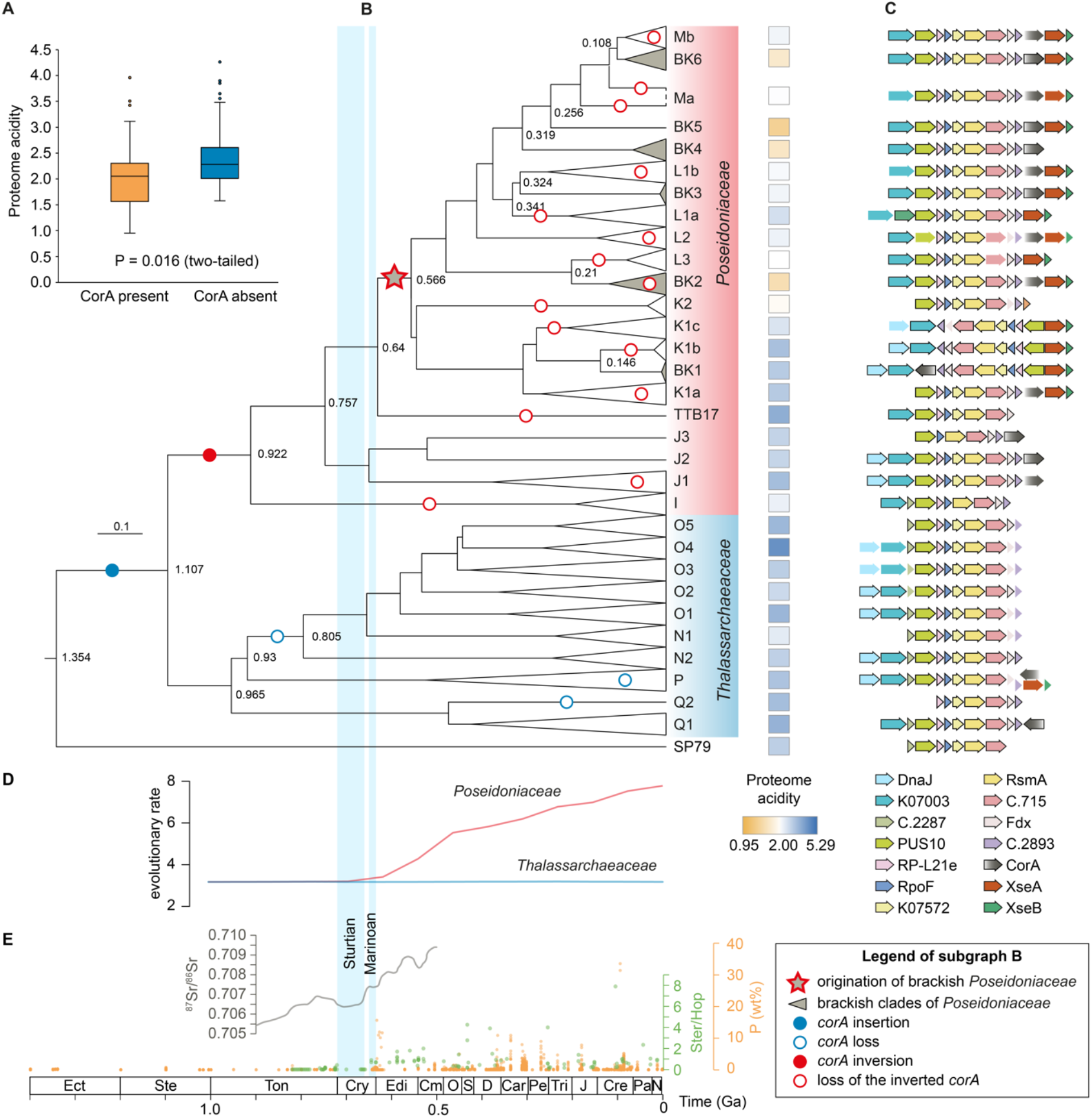
Genetic changes, evolutionary rate and geological background in the origination and evolution of brackish *Poseidoniales*. (**A**) The distribution of proteome acidity in genomes with or without the *corA* gene. (**B**) Molecular dating results and proteome acidity patterns of *Poseidoniales* subclades with the change of the *corA* gene in evolution. The tree is part of the tree in fig. S6A. Proteome acidity levels at the right show the median values of MAGs in each clade. (**C**) Arrangement of the stress-response gene cluster in *Poseidoniales* genomes. Arrows without edge suggest that the genes are present only in some of the MAGs in each clade. Potential functions of genes are explained in Supplementary Text. (**D**) Evolutionary rates of *Poseidoniaceae* and *Thalassarchaeaceae*. (**E**) Geological records of P deposit in shales from Reinhard *et al*. 2017 (Reinhard et al., 2017), the ^87^Sr/^86^Sr curve copied from Fig. 1 in Laakso *et al*. 2020 (Thomas A. Laakso et al., 2020), and the relative contribution of eukaryotic and bacterial lipids to sedimentary organic matter approximated by the sterane/hopane ratio (Ster/Hop) by Brocks *et al*. (Brocks et al., 2017). Glaciation events are based on Hoffman *et al*. 1998 (Hoffman et al., 1998). Ect = Ectasian, Ste = Stenian, Ton = Tonian, Cry = Cryogenian, Edi = Ediacaran, Cm = Cambrian, O = Ordovician, S = Silurian, D = Devonian, Car = Carboniferous, Pe = Permian, Tri = Triassic, J = Jurassic, Cre = Cretaceous, Pa = Paleogene, N = Neogene. Ga = billion years ago.

Neighboring gene analysis integrated with phylogenetic interference analysis indicates that *corA* had a highly conserved evolutionary trajectory and may have played an essential role in stress response. Specifically, the tree of *corA* is highly congruent to the phylogenomic tree of *Poseidoniales* and the adjacent Marine Group III (MGIII) archaea suggesting that after its gain in the common ancestor of *Poseidoniales*, *corA* was generally passed vertically when *Poseidoniales* diversified (Fig. 2B, fig. S5 and Supplementary Text). The absence of *corA* the majority of *Thalassarchaeaceae* and in some *Poseidoniaceae* is likely a result of sporadic loss (fig. S4 and Supplementary Text). Moreover, *corA* is exclusively found at the tail of a highly conservative gene cluster consisting of over ten syntenic genes (Fig. 2C and fig. S4). This gene cluster contains core gene sets involved in DNA repair, transcription, translational regulation, and post-translational modification by modulating macromolecules such as DNAs, RNAs and proteins (Supplementary Text). As adjacent and syntenic genes often form operons and transcribe simultaneously (Gao et al., 2020), genes in this cluster may be regulated in concert with each other in stress response.

Importantly, we find the sporadic distributed of brackish clades in the *Poseidoniaceae* evolutionary tree had a single origination possibly resulting from the inversion of *corA* in the common ancestor of *Poseidoniaceae* as in the two basal genera of *Thalassarchaeaceae* P and Q1, *corA* genes are in reverse coding direction of those in *Poseidoniaceae* (Fig. 2BC and see discussion in Supplementary Text). Consequently, *corA* was in the same coding direction as the rest of the gene cluster making co-transcription possible. However, after this inversion event, *corA* did not seem to immediately establish coordination with adjacent genes in stress-response since a basal monophyletic clade of *Poseidoniaceae* consisting of genera J1, J2 and J3 partially contained *corA* but did not show typical brackish adaptation features. Instead, the origination of brackish *Poseidoniaceae* likely only occurred in the common ancestor of genera K1, K2, L1, L2, L3 and M, in which *corA* might have fully coupled its regulation and transcription to the gene cluster (Supplementary Text). This rearrangement provided a selective advantage for *Poseidoniaceae* in adaptation to brackish waters and *corA* was thus preserved in further diversification. The following sporadic loss of *corA* gene in other *Poseidoniaceae* lineages, however, may force them to go back to the saline water (Fig. 2B).

### A proposal of two step changes in habitat transition by *Poseidoniaceae*

Genetic mechanisms are elusive in the study of evolutionary transitions across the salinity barrier. Eiler *et al*. suggested a gradual tuning of metabolic pathways and transporters of marine microbial clades towards organic substrates in freshwater environments (Eiler et al., 2016). In contrast, Henson *et al*. proposed that the irreversible loss of osmolyte transporters could be the critical step in the formation of the freshwater bacterial clade (Henson et al., 2018). Here we use the sporadic loss of the *corA* gene in *Poseidoniaceae* genera to elucidate the evolutionary history of *Poseidoniaceae* transitioning between brackish and marine habitats, which is characterized by at least ten salinity-based divergent events (Fig. 2A). As an example, we tracked the gene-gain and gene-loss events in association with the salinity and proteome acidification diversification in further selected dataset of high-quality MAGs (completeness median = 93.63%, contamination median = 0, table S2) in subgroups BK4, BK5, BK6 and M based on the ALE results (table S3).

A two-step genomic change was identified during the marine-brackish inter-transition (Fig. 3). The first step was the gain or loss of the *corA* gene. Specifically, the gene was replaced by a new copy in the common ancestor of BK4 and then vertically transferred in this subgroup. In the monophyletic clade containing BK5, M and BK6, *corA* was vertically transferred followed by sporadic losses in marine species of the M subgroup (Fig. 3).

**Fig. 3.**
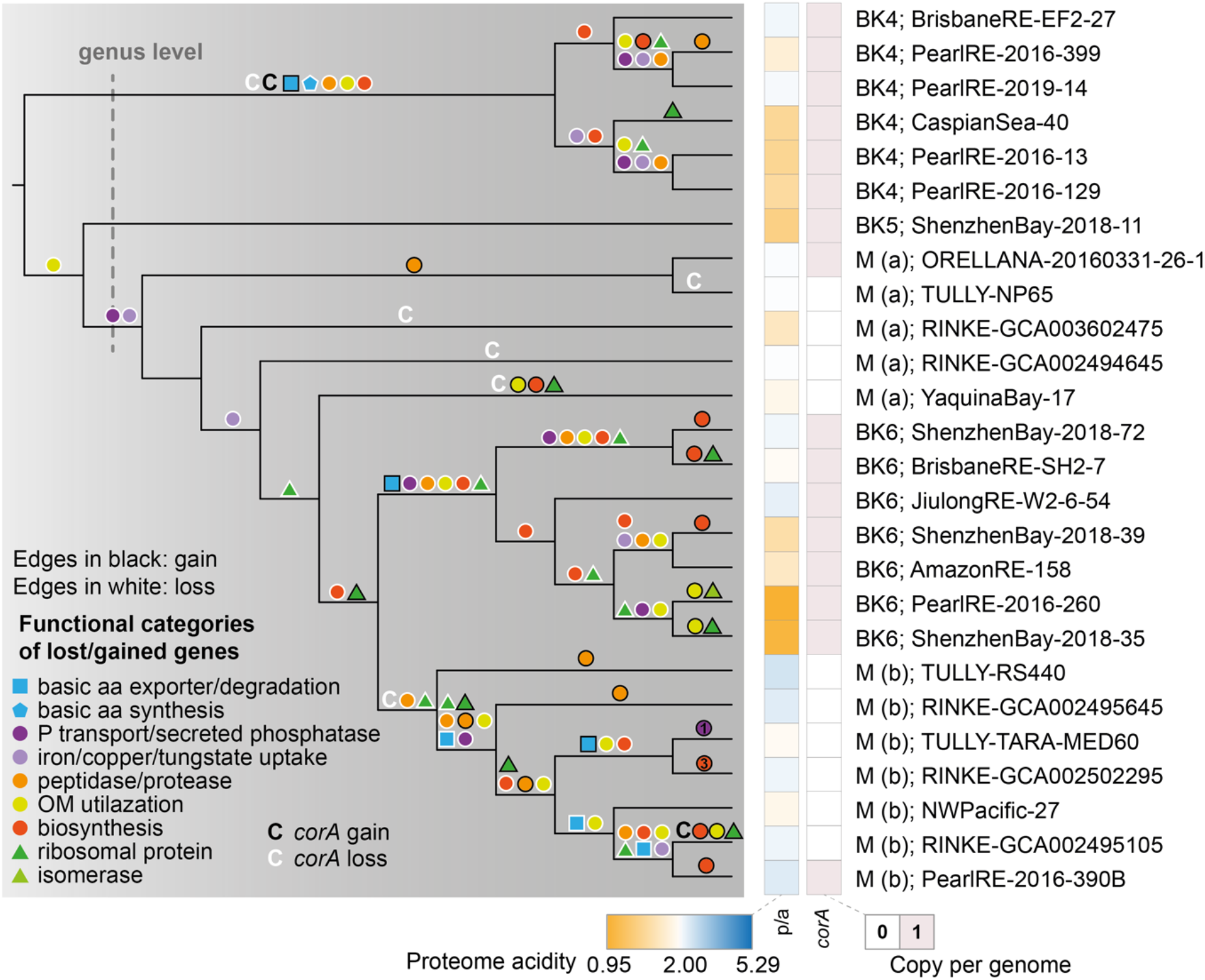
Essential gene-gain and gene-loss events in the marine-brackish divergence of the BK4-BK5-M-BK6 monophyletic clade of *Poseidoniaceae*. The cladogram of the BK4-BK5-M-BK6 clade of fig. S7A is shown. Genus-level cutoff in the tree is according to GTDB taxonomic ranking. Gene-gain and gene-loss events between adjacent internal nodes or between adjacent internal nodes and terminal taxa are illustrated on relevant branches. Capital C means the *corA* gene. Shapes with black edges or letters in black show gene gain events, while that with white edges or letters in white show gene loss events.

In the second step, accompanying and especially following the change of *corA*, massive gains and losses of habitat-specification genes possibly mediated by LGT are observed in *Poseidoniaceae* genomes during their gradual diversification to marine or brackish environments, respectively (Fig. 3). For example, as the less acidified proteomes of brackish *Poseidoniaceae* comprise more basic amino acids than that of their marine counterparts (fig. S1B), the concentration of free basic amino acids in the cytoplasmic pool needs to be lowered by either increasing export or stopping biosynthesis to maintain charge balance. Indeed, the acquisition of lysine/arginine efflux and the loss of lysine biosynthesis happened in the ancestors of BK4 and BK6, respectively (Fig. 3). Moreover, microorganisms living in brackish and marine environments face great physiochemical differences in nutrient availability, substrate composition and stress types. Accordingly, we find large-scale loss of genes involved in phosphorus and low-abundant metal uptake, peptide degradation (peptidases/proteases), and organic matter utilization in BK4 and BK6 lineages, reflecting a relative eutrophic brackish environment where extra substrate transport machinery is unnecessary. At the same time, some peptidases and biosynthetic enzymes required for brackish water adaptation were obtained in brackish taxa. For example, the acquisition of fructose/tagatose bisphosphate aldolase by genome ShenzhenBay-2018-35 and 2-keto-4-pentenoate hydratase in the catechol meta-cleavage pathway by genome PearlRE-2016-260 possibly reflects the adaptation of brackish *Poseidoniaceae* to consume terrestrial substrates such as lignin (Bugg et al., 2011) (table S3).

This observation is consistent with the fundamental rule that salinity is the strongest environmental barrier in microbial habitat transition and highlights a two-rank structure of selection and genetic change in microbial transition between ecologically distinct environments. Specifically, the sudden change of primary and qualitative niche trait (e.g. adaptation to salinity fluctuation in this study) precedes the gradual changes in secondary and accumulative traits (e.g. proteome acidity change and multidimensional metabolism adjustment) in *Poseidoniaceae* genomes.

### Habitat divergence and evolutionary rate changes of *Poseidoniales* potentially affected by the enhanced land weathering and the rise of algae after the snowball Earth

Temporal connection of genetic changes with major geological events can provide insights into the potential environmental driving force of microbial evolution in the absence of a fossil record. Despite the frequent discussion on uncertainty of methodology that may impair dating accuracy, molecular clock analysis aided by biomarker records has unveiled important events on life evolution before the emergence of multicellular organisms (Betts et al., 2018; Blank, 2009; Boden et al., 2021; Chen et al., 2020; Ngugi et al., 2023; Ren et al., 2019; Yang et al., 2021). We here establish a geological time scheme in the evolutionary diversification of *Poseidoniales* to infer the possible geological and environmental driving force in the origination of brackish lineages (Fig. 2B). To ensure the robustness of molecular dating analysis on *Poseidoniales*, we here compare the results from alternative phylogenetic tree topologies (i.e. ‘methanogen-basal’ v.s. ‘DPANN-basal’, see Supplementary Text), select a modeling method based on the justification in our previous study (Yang et al., 2021), and build our conclusions on multiple interconnected lines of genetic and geological evidence (Fig. 2).

Our dating analysis shows that the origination of brackish *Poseidoniaceae* was temporally linked to the emerge of eukaryotic algae in the Ediacaran Period. Based on the ‘methanogen-basal’ tree topology, the emergence of *Poseidoniales* and divide of *Poseidoniaceae* and *Thalassarchaeaceae* happened around 1.107 billion years ago (Ga) (Fig. 2B, fig. S6A and Supplementary Text). The divergence of extant genera of *Poseidoniaceae* and *Thalassarchaeaceae* happened before 0.9 Ga. The insertion of *corA* to the common ancestor of *Poseidoniales* was in the period from 1.354 to 1.107 Ga and its inversion happened between 1.107 and 0.922 Ga (Fig. 2B). Notably, the latest common ancestor of extant brackish *Poseidoniaceae* (*i.e.* the latest common ancestor of genera K1, K2, L1, L2, L3 and M as we have mentioned above) likely emerged between 640 and 566 Ma (561 and 496 Ma according to the ‘DPANN-basal’ tree, Fig. 2B and fig. S6B), a period right after the Marinoan Glaciation (approximately 650 to 632.3 Ma) and covering most of the Ediacaran period. This period is also highlighted by the rise of eukaryotic algae (Brocks et al., 2017), which might have enhanced the production of organic substrates for the growth of *Poseidoniaceae* (Zhang et al., 2015). After that, the divergence of brackish and marine lineages of *Poseidoniaceae* happened multiple times in an algal-dominant ocean until a most recent major divide between the subclades BK6 and Mb in 108 Ma (Fig. 2B).

The evolutionary rates of *Poseidoniaceae* and *Thalassarchaeaceae* were remarkably different in the past 700 Ma, probably reflecting different levels of influence from the land (Fig. 2D). The rapid increase in evolutionary rate of *Poseidoniaceae* started generally from the Cryogenian glaciation and accelerated in the Ediacaran-Cambrian period. After that, the evolutionary rate increased in a constant pattern till present. This tendency is consistent with the increase of continental weathering (Halverson et al., 2010; Sharoni and Halevy, 2023), coastal sedimentary phosphorite (Reinhard et al., 2017), and algae rise (Brocks et al., 2017). It is also consistent to the origination and diversification of brackish species *Poseidoniaceae* after the Snowball Earth and throughout the Phanerozoic (Fig. 2E), implying a potential impact of land nutrient supply and consequently coastal algal bloom on the evolution of nearshore inhabiting *Poseidoniaceae*. In contrast, the evolutionary rate of *Thalassarchaeaceae* stayed at a low level in the past one billion years (Fig. 2D) in line with their preferred environments that are oligotrophic and distant from the shore.

## Conclusion

Origination of global brackish microorganisms is a long-standing question. In this study, we apply detailed genome comparison and evolutionary analysis to investigate the key and detailed genetic changes causing the origination of the brackish lineages of the most abundant archaeal heteroplankton *Poseidoniales*. We find that the divergent and globally distributed brackish lineages of *Poseidoniaceae* have a single origination. Importantly, we identify the *corA* gene to be the key gene for adaptation in brackish environments, which may regulate intracellular magnesium concentration to stabilize macromolecules. The regulation of this gene was likely coupled to a highly conserved archaeal stress-response gene cluster after an ancient inversion event allowing concerted responses to multiple stresses including fluctuating salinities in brackish waters. Importantly, we find that the losses and gains of *corA* were followed by metabolic acclimation and diversification mediated by lateral gene transfer of substrate-specific genes. Molecular dating and geological inference suggest brackish *Poseidoniaceae* in Ediacaran-Cambrian possibly flourished due to the rise of algal productivity in coastal and even brackish waters because of enhanced phosphorus supply in response to land weathering (Dodd et al., 2023).

Our results highlight coordination of genome rearrangement and global environmental changes driven by major geological events in the emergence of novel groups of planktonic archaea. On the other hand, the emergence of these brackish microbes filled the new niche gaps in the Ediacaran-Cambrian coasts and contributed to the recycling of organic matter produced by algae through the microbial loop and may have been an important player of microbial carbon pump in that time (Jiao et al., 2010; Zhang et al., 2018). The accelerated evolutionary rate of coastal *Poseidoniales* since the Ediacaran period indicates that evolution of both the macroheterotrophs (the animals) (Erwin et al., 2011) and microheterotrophs (*Poseidoniaceae* here and possibly some heterotrophic bacteria) were stimulated since late-Proterozoic. Therefore, these flourished heterotrophic archaea and bacteria could have given a previously overlooked contribution to the biomineralization of organic matter in seawater (Peters and Gaines, 2012) and consequently further accelerated land weathering after the Snowball Earth.

We are, however, aware of the caveats that are inherited with molecular clock calculation of geological ages of ancient microorganisms, which are improving in precision given the growing database of high-quality genomes and rapid development in methodology (Ren et al., 2019; Szöllõsi et al., 2022; Wolfe and Fournier, 2018). We expect that our prediction of the genetic changes in habitat specificity of *Poseidoniales* can be further validated with *in vivo* experiment when pure or enriched cultures are available (Zhang et al., 2015). In addition, modeling Ediacaran coastal hydrodynamic and biogeochemical cycles with sedimental records of biomarkers for specific brackish microorganisms may provide further support to our findings.

### A model integrating genetic and geological explanations attests to the origination of brackish

#### Poseidoniaceae

The emergence of novel life forms in earth’s history was likely trigged by a mixture of geological and ecological changes while genetic innovation made it possible. However, collecting mutually corroborated lines of evidence to prove such ancient processes is a formidable task especially for microorganisms that lack fossil records. Here, based on the detailed genetic features of brackish versus marine lineages and the trend of global environmental changes on continental margins in history, we reconstruct the possible trajectory in the origination of brackish *Poseidoniaceae* (Fig. 4). After the divergence of *Poseidoniaceae* and *Thalassarchaeaceae* in about 1.1 Ga, the *corA* gene inserted to the end of an archaeal stress-response gene cluster in earlier time was inverted providing a genetic preparation for brackish adaptation. About 300 million years later in the postglacial Ediacaran coasts, rapid increase in phosphorus from land supply and organic-matter release might have led to the rise of algae. The nutrient-stimulated algal boom should be most significant in coastal waters and the flourishing of algal-derived organic matter then fed the emerging animal species (Knoll, 2017) in a transforming food web on continental shelf. These algal-derived organic matter also accelerated the evolution and diversification of coast-inhabiting *Poseidoniaceae*, possibly along with other heterotrophic bacteria and their potential predators – protists or filter-feeders – and resulted in a highly competitive community in the Ediacaran-Cambrian coasts. At the same time, enhanced nutrient supply might have stimulated algal bloom in the expanded brackish water bodies during the postglacial transgression (Hurtgen et al., 2006). This food-rich but predator-less environment formed a remarkable niche gap. Consequently, *Poseidoniaceae* populations were continually selected when (1) the *corA* gene was in better coordination in transcription and regulation with the stress-response gene cluster to tolerant the stress of salinity fluctuation, (2) the proteome was less acidified to meet lower average salinity, and (3) the metabolism was better adapted to more eutrophic environments. This gradual tuning in gene regulation, proteome amino acid composition and substrate metabolism finally led to the origination of brackish *Poseidoniaceae*. Later, when land-weathering and phosphorus supply kept increasing in Phanerozoic, some brackish lineages continued exploring upstream habitats with salinity down to 1‰ and even developed metabolism to utilize land-plant derived organic compounds, which increased rapidly from Silurian to Carboniferous (T.A. Laakso et al., 2020). On the other hand, some lineages lost the *corA* gene and returned to marine habitats. In some rare cases, they might further gain the *corA* genes possibly through recombination and shuffle between marine and brackish niches in expanded shallow marine environments (Peters and Gaines, 2012).

**Fig. 4.**
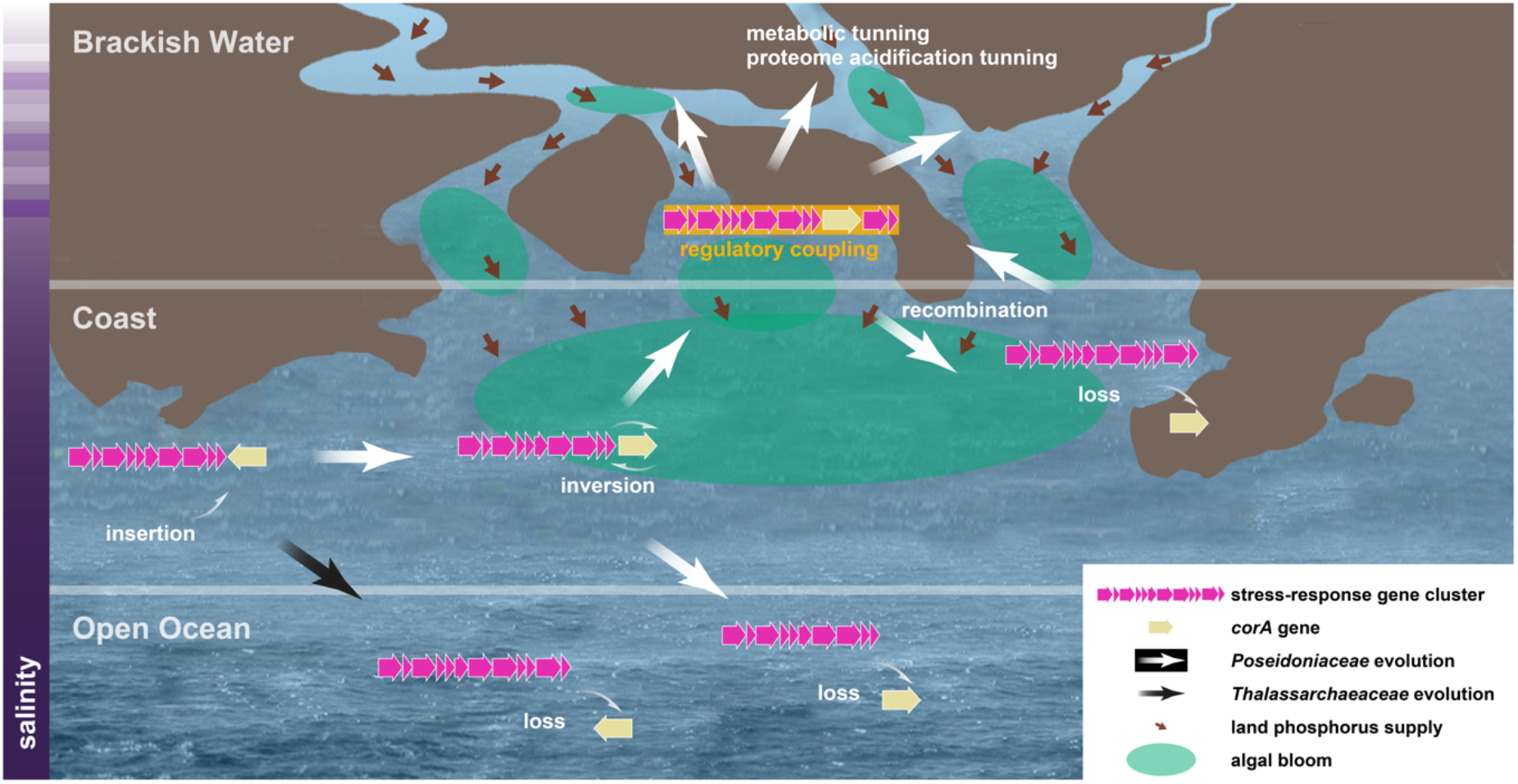
A schematic diagram shows the process of genetic changes during the transition of *Poseidoniales* between salinity distinct habitats and environmental features in the origination of brackish *Poseidoniaceae* in the Ediacaran-Cambrian coasts.

## Methods

Microplanktons were collected at the Pearl River estuary in 2011, 2012, 2016 (Xu et al., 2022), 2018 (in the adjacent Shenzhen Bay) and 2019 (National Omics Data Encyclopedia (NODE, https://www.biosino.org/node/) project: OEP001662), the Jiulong River estuary in 2018 (NODE project: OEP000961), the Yangtze River estuary in 2016 (NODE project: OEP001524), the Brisbane River estuary in 2020 (NCBI project: PRJNA872317), and the Northwest Pacific in 2015 and 2017 (NODE project: OEP001662) (table S1). Immediately after collection, surface water was first filtered through 2.7 μm pore-size glass fiber filters (Shanghai Mosutech, Shanghai, China) to remove large particles and the filtrates were then filtered through 0.22 μm pore-size membrane filters (Pellicon cartridge, Millipore Corp., Billerica, MA, USA) to collect microbial cells. Filters were then frozen in liquid nitrogen and stored at −80°C in lab till further processes. DNA was extracted by using the FastDNA SPIN kit for soil (MP Biomedicals, Solon, OH, USA) following the manufacturers’ instructions. Metagenome sequencing was conducted on an Illumina HiSeq 2500 platform at Novogene Bioinformatics Technology Co., Ltd. (Beijing, China). Raw reads of the published metagenomes of the Caspian Sea (Mehrshad et al., 2016), the Baltic Sea (Alneberg et al., 2020, 2018; Hugerth et al., 2015; Larsson et al., 2014), the Columbia River estuary (Fortunato and Crump, 2015), the Amazon River estuary (Satinsky et al., 2014), the Yaquina Bay estuary (Kieft et al., 2018), the Helgoland coast (Orellana et al., 2019), the Port Hacking coast (Rinke et al., 2019), and the Tara Oceans Project (Pesant et al., 2015) were downloaded from public databases (table S1). Clean reads of the above metagenomes were generated by using the reads_qc module of MetaWRAP (v. 1.2.1) (Uritskiy et al., 2018).

Description of the generation of the non-redundant *Poseidoniales* genome dataset, phylogenomic analysis, MAG abundance calculation and functional annotation, proteome acidity and habitat salinity analysis, *Poseidoniales* evolutionary analysis including gene gain/loss tracking, and the molecular clock analysis can be found in Supplementary Methods.

## Acknowledgements

We thank Zongjun Yin (Nanjing Institute of Geology and Palaeontology, Chinese Academy of Sciences) and Weiqi Yao (Southern University of Science and Technology) for their valuable suggestions in geological records. Computation in this study was supported by the Centre for Computational Science and Engineering at the Southern University of Science and Technology.

## Author contributions

Lu Fan and Chuanlun Zhang conceived this study. Bu Xu, Songze Chen, Fuyan Li, Wei Xie, Apoorva Prabhu, Dayu Zou, Ru Wan, Hongliang Li, Haodong Liu, Yuhang Liu, Shuh-Ji Kao, Jianfang Chen, Yuanqing Zhu, Christian Rinke, and Meng Li collected the samples and extracted DNA. Lu Fan, Bu Xu, Songze Chen, Yang Liu, Fuyan Li, Apoorva Prabhu, Dayu Zou, Ru Wan, and Hongliang Li analyzed the metagenome data, produced the genomes, and conducted all other analyses. Lu Fan, Bu Xu, Maoyan Zhu, and Chuanlun Zhang interpreted the results and drafted the manuscript. All authors contributed to the final version of the manuscript.

## Additional Information

Supplementary Information is available for this paper.

