## Supplementary Information for "Gene inversion triggered origination of brackish archaeal heterotrophs in the aftermath of the Cryogenian Snowball Earth"

### Supplementary Text

#### 1. Acidification of *Poseidoniales* proteomes

The distribution of protein isoelectric points in *Poseidoniales* proteomes showed an acidic bias similar to what was observed in halophiles but in less extent [1] (fig. S8). Specifically, there are two major bumpy zones, one generally peaks at p*I* 4.6 (named 'acidic peak') and the other at p*I* 6.25 (named 'semi-acidic peak'). The alkaline peak usually found around p*I* 10 in other marine bacteria and archaea [2] is hardly detectable. Therefore, *Poseidoniales* likely have high intracellular ion concentration [3]. Brackish *Poseidoniaceae* usually have lower acidic peaks but higher semi-acidic peaks than their marine counterparts, suggesting that less acidified proteomes are associated with lower habitat salinity [2].

#### 2. The distribution of osmotic regulation genes in *Poseidoniales* genera

The regulation of cellular osmotic pressure of a microbial cell is often accomplished by coordinated multiple ions and small molecule channels or transporters [4]. Besides *corA* and *znuABC*, we conducted a comprehensive search for ion channels, ion transporters, osmolyte transporters, osmolyte synthetases, and other proteins potentially related to osmotic regulation according to the KEGG, arCOG, and COG databases.

Ten additional types of relevant proteins or protein complexes were found in more than three genera of *Poseidoniales* (fig. S4) and seven of them were present in nearly all *Poseidoniales* genera. The ubiquitous distribution of TrkH, PutP, HppA, NhaD, KhtT, ChaA, YrbG, and MscS genes suggests that *Poseidoniales* may employ multiple strategies to cope with extracellular salinity change in seawater.

1) The p*I* distribution of *Poseidoniales* proteomes shows typical acidification, suggesting that they may import cations to balance osmotic pressure in seawater (fig. S8). Indeed, potassium transporter Trk is present in all *Poseidoniales* genomes (fig. S4). TrkH is an ATP-activated potassium transport protein [5]. Like many halophilic bacteria and archaea, *Poseidoniales* cells may actively and selectively import potassium ions and maintain a high concentration of potassium ions to keep osmotic balance in seawater. This hypothesis is consistent with the acidified proteomes observed in halophiles [6] (fig. S8).

2) Microorganisms may import or synthesize compatible solutes as well as ions in response to osmotic stress [7]. Proline is one of such frequently used organic osmolytes. *Poseidoniales* encode PutP/NPSP, a $Na^+$:proline symporter, and may be able to conduct sodium-dependent uptake of extracellular L-proline [8].

3) *Poseidoniales* cells may directly extrude sodium and potassium ions. They commonly encode a pyrophosphate-energized sodium pump HppA [9], a $Na^+/H^+$ antiport NhaD [10], a $Na^+/H^+$ antiport KhtT [11], a non-specific $Ca^{2+}$,$Na^+$, $K^+/H^+$ antiport ChaA [12,13] and a $Na^+/Ca^{2+}$ exchanger YrbG [14]. They also encode the *mscS* gene, a mechanosensitive channel releasing solutes in speedy and non-discriminating manners [15].

An extensive analysis of genes potentially involved in osmotic regulation suggests that three proteins and protein complexes are possibly responsible for niche differentiation of *Poseidoniaceae* and *Thalassarchaeaceae* [16] (Fig. 2C, fig. S4). The three genes including *natB*, *kefB* and *mgtA* were present in all *Poseidoniaceae* genera but only in some evolutionary basal *Thalassarchaeaceae* genera (Fig. 2C, fig. S4). NatB is the permease component of NatAB, an ABC-type transport system that extrudes intracellular sodium ions [17]. KefB belongs to the glutathione-gated potassium efflux system [18]. MgtA is involved in ATP-activated $Mg^{2+}$ import. It is induced at very low external $Mg^{2+}$ concentrations [19].

As coastal surface water is frequently diluted by land-input fresh water, coastal microplanktons may experience sudden drop of extracellular salinity. Having additional $Na^+$ and $K^+$ exporters can be a great advance for cells to maintain osmotic balance against such a stress. Moreover, magnesium imported through MgtA may stabilize proteins and RNAs in additional to basic physiological demands of magnesium in occasionally diluted coastal water, which lacks magnesium (see below). *Poseidoniaceae* are found to be dominant in coastal areas where algal oligosaccharides would be more readily available, while most *Thalassarchaeaceae* are adapted to mesopelagic and oligotrophic waters where direct algal inputs are limited [16,20]. The skewed distribution of these three genes may explain the observation that *Poseidoniaceae* are more adapted to dynamic coastal waters while *Thalassarchaeaceae* generally restrict their habitat in pelagic zones [16].

**3. Putative functions of the genes in the syntenic cluster**

The *corA* gene is inserted in a cluster of thirteen syntenic genes (Fig. 2B, fig. S4). This gene cluster is conserved in all *Poseidoniaceae* and *Thalassarchaeaceae* genera and was possibly inherited from the common ancestor of MGII. Among these genes, chaperone protein DnaJ can be involved in stress responses [21]. Homologs of DnaJ in plants were found to enhance salinity tolerance [22]. The protein annotated as K07003 is a resistance-nodulation-division transporter possibly functioning as a drug efflux pump in detoxification [23]. tRNA pseudouridine synthase 10 (PUS10) is only found in archaea and eukaryotes. It is involved in post-transcriptional modification by tRNA modification [24] and is salinity responsible [25]. Large subunit (50S) ribosomal protein L21e (RP-L21e) catalyzes the peptidyl-transfer reaction of messenger RNA-directed protein bio-synthesis [26]. In soybean, this protein is downregulated under short-term of salt stress [26], but upregulated in long-term of salt exposure [27]. RpoF is the F subunit of an archaeal DNA-directed RNA polymerase [28] and is found responsible in heat tolerance [29]. The protein annotated as K07572 contains an RNA-binding PUA domain. Its homolog in found functioning in translational regulation by modulating mRNA in human cells [30]. Ribosomal RNA small subunit methyltransferase A (RsmA/KsgA) modify rRNAs [31]. Ribosome methylation can facilitate selective translation in response to environmental stresses [32]. PsiBLAST search for the protein C.715 suggests that its sequence is distantly related to DNA-directed RNA polymerase subunit A of *Candidatus* Syntrophoarchaeum. The protein Fdx is likely a ferredoxin. However, its exact function is unknown [33]. XseA and XseB are the large and small subunits of exonuclease VII, respectively. Exonuclease VII is involved in DNA repair and recombination that hydrolyses single-stranded DNA from both 5' and 3' ends [34]. Proteins C.2287 and C.2893 are unclassified.

In summary, among the eleven annotated genes, two served in DNA protection (XseA and XseB), two in transcription (RpoF and C.715), and four in translational regulation likely by tRNA and ribosome modification (PUS10, RP-L21e, K07572 and RsmA) and one in post-translational modification (DnaJ). Therefore, genes in this cluster likely function as regulatory hubs in the metabolic networks of *Poseidoniales* in stress response. The highly conservative association and organization of these essential genes in *Poseidoniales* genomes suggest that they may be coordinated as a cohort and by unknown universal regulatory factors to environmental stresses as an analogy to microbial immune components [35].

**4. Possible functions of CorA and ZnuABC in *Poseidoniales***

At least 133 of the 159 *Poseidoniales* MAGs (83.6%) in fig. S4 encode either *corA*, *znuABC*, or both, suggesting that the import of divalent cations may be crucial to the survival of *Poseidoniales*. CorA is one of the main import channel of magnesium ions ($Mg^{2+}$) in bacteria and archaea[36] and is also capable of transporting similar divalent ions such as $Co^{2+}$ and $Ni^{2+}$ and $Zn^{2+}$ [37]. In addition, gated transport of CorA is regulated by intracellular $Mg^{2+}$ level [38] and CorA may also export $Mg^{2+}$ [37].

Magnesium is the most abundant divalent cation inside the cell. It is mainly bound to nucleic acids, negatively charged phospholipids, and proteins [39]. It is an essential cofactor for numerous enzymatic and metabolic pathways [40]. To facilitate the adaptation of brackish *Poseidoniaceae*, cation magnesium may act as a DNA compaction agent. It can effectively bind to the chromosome, similar as histones or other positive charged proteins [41], to stabilize the chromosome [42], facilitate genome-wide DNA compaction and inhibit transcription [43]. We did not find classical archaeal histone sequences in *Poseidoniales* genomes. Therefore, *Poseidoniales* may partly depend on divalent cations such as $Mg^{2+}$, $Zn^{2+}$, or $Mn^{2+}$ in DNA compaction. At osmotic stress, the import of additional $Mg^{2+}$ may facilitate DNA compaction and reduce gene transcription of brackish *Poseidoniaceae* cells. Moreover, in dynamic estuarine environments, salinity fluctuation leads to sudden change in intracellular ion levels causing strong detrimental effects on the structural stability of structural or catalytical macromolecules such as proteins and RNAs. Import of $Mg^{2+}$ through CorA may bind to the exposed negative groups and stabilize these macromolecules.

ZnuABC is an ABC-type, high-affinity $Zn^{2+}$-specific uptake system [44]. Homologs of ZnuABC may also import manganese [45]. Zinc serves in structural and catalytic roles including enzyme cofactor, stabilizing protein structure [46]. It can modulate ribosomal proteins [47] and DNA-binding proteins [48]. Manganese is involved oxidative stress resistance [49]. It can also interact with chemical groups in DNA and proteins stimulating DNA condensation[50]. Therefore, the imported zinc or manganese through ZnuABC may also serve to stabilize macromolecules or facilitate DNA compaction as magnesium may do.

**5. Evolutionary trajectory of the *corA* gene and the *znuABC* gene set in *Poseidoniales***

The *corA* gene is found in some Marine Group III (MGIII) Euryarchaea genomes but not in the conserved stress-response gene cluster (fig. S4). In the partial SP9 genome, which branched before the divergence of *Poseidoniaceae* and *Thalassarchaeaceae*, the stress-response gene cluster is detected but the *corA* gene is absent (Fig. 2B, fig. S4). Therefore, the *corA* gene was inserted into this highly conserved stress response gene cluster in one of the common ancestors of *Poseidoniales*. Notably, in the three genomes of basal *Thalassarchaeaceae*, *corA* was in opposite coding direction to the rest of the gene cluster while in *Poseidoniaceae* it was always in the same coding direction. Such an arrangement suggests an inversion event in the ancestor of either *Poseidoniaceae* or *Thalassarchaeaceae*. However, it is unknown whether *corA* was initially inserted in the same or opposite direction to the rest genes of the cluster. A likely scenario is that in the common ancestor of *Poseidoniales*, the *corA* gene was inserted in opposite direction to the gene cluster. It was then lost during the early diversification of *Thalassarchaeaceae* since this insertion would disrupt the integrity of the stress response gene cluster and might be deleterious, at least partially. This assumption can explain that only three copies of *corA* in *Thalassarchaeaceae* are found in opposite direction to the gene cluster in the two basal genera (Fig. 2B and fig. S4).

Notably, in basal genera of *Poseidoniaceae* including J1, J2 and J3, *corA* is not direct adjacent to *xseAB*, suggesting the regulation of *corA*, *xseAB* and other upstream genes may not yet be fully coordinated (Fig. 2B, fig. S4). In genus K1, part of the gene cluster including *corA* and other eight genes were inverted to the rest of the genome. These variations imply that the stress-response gene cluster including *corA* was in progressive evolution in regulatory coordination until genus L3/BK2 and afterward.

In addition, we generally observe a mutually exclusive distribution of the zinc/manganese ABC transport complex gene set *znuABC* with *corA* in *Poseidoniaceae* MAGs (Fig. S4). The *znuABC* gene set is present in 6.1% MAGs (2/33) of the brackish clades and in 67.2% MAGs (248/369) of the marine clades. This pattern suggests that these two ion transporters may be functionally redundant as both magnesium and zinc are divalent cations and may aid in stability of intracellular macromolecules (fig. S4 and Supplementary Text). In contrast to *corA*, the evolution of *znuA* in *Poseidoniales* was often mediated by lateral gene transfer (LGT) (fig. S5) including 39 gain and 35 loss events as predicted by amalgamated likelihood estimation (ALE) (fig. S4). Moreover, the *znuABC* gene set is not associated with any specific genes or gene clusters (fig. S4). This finding supports a genome evolutionary model that the potential regulatory coupling of magnesium import by CorA with other essential stress-response functions was the strongest determinant of brackish *Poseidoniaceae*, whereas in marine *Poseidoniaceae* whose *corA* was lost from the stress-response gene cluster, *znuABC* was then obtained as compensation for divalent cation transporter. Notably, 37 of the 42 contigs (88.1%) encoding the *corA* gene and 114 of the 122 contigs (93.4%) encoding the *znuA/B/C* gene(s) contain at least five ORFs (Fig. S4). These long contigs have sufficient information of read coverage and genome composition features and thus were unlikely to be falsely binned by our binning and decontamination approaches (Supplementary Methods), deeming the presence/absence analyses of *corA* and *znuA/B/C* genes to be credible.

**6. Alternative rooting in the tree of Archaea**

To conduct the molecular clock analysis of the evolution of *Poseidoniales*, we first attempted to place *Poseidoniales* in the tree of Archaea. Previous studies assigned *Poseidoniales* and MGIII Euryarchaea as sisters of [51] or as part of [52–54] *Thermoplasmatota*. To elucidate the accurate phylogenetic relationship of *Poseidoniales* with basal *Thermoplasmatota* lineages, we downloaded genomes belonging to *Thermoplasmatota* from GTDB [54] and NCBI [55] and generated a non-redundant genome dataset containing 188 genomes including *Thermoplasmatota*, other archaea plus the outgroup (table S2 and table S4). We then conducted phylogenetic analysis based on 39 selected marker genes. To examine potential artifacts caused by compositional heterogeneity, we removed 5%, 10%, 20%, 40% and 60% of the most heterogeneous sites in the alignment matrix (fig. S7). *Poseidoniales* and MGIII archaea form a monophyletic clade with EX4484-6 (a class-level group) of

*Thermoplasmatota*. This clade is in sister relationship with the class *Thermoplasmata* (fig. S7). Such a topology is stable but until 20%, 40% or 60% of heterogeneous sites are removed, when EX4484-6 and *Thermoplasmata* form a monophyletic clade which is in sister relationship with the clade of *Poseidoniales* and MGIII archaea (fig. S7).

Based on this phylogenetic frame of *Thermoplasmatota*, we further built a dataset containing 230 taxa including 165 representative *Poseidoniales* taxa, representative genomes of other *Thermoplasmatota* major clades, representatives of *Halobacteriota*, *Thermoproteota*, *Methanobacteriota*, and I*ainarchaeota*. Taxa belonging to *Thermoplasmataceae*, *Sulfolobaceae*, *Thermofilaceae*, *Thermocladiaceae*, and *Thermoproteaceae* were included for downstream time calibration (fig. S7). However, tree topology examinations by removing certain proportion of compositional heterogeneous sites generated two conflicting structures at the root of Archaea: 'methanogen-basal' (fig. S9ABC) and 'DPANN-basal' (fig. S9DEF). This conflict was frequently debated in recent studies [52,54,56,57]. Importantly, we carefully justified the fundamental impact of basal-branch topology of the Archaea tree on molecular clock analysis. Although currently the 'DPANN-basal' topology is favored over the 'methanogen-basal' one, a consensus is yet to reach by the research community. As the root of archaea is a key constraint for time calibration [58], it is necessary to justify the impact of alternative tree topology on the time estimation of *Poseidoniales* evolution. Phylogenetic trees generated from the untreated, 5%-, 10%-, 20%-, 40%- and 60% removal of heterogeneous sites produced two distinct topologies of the *Iainarchaeota* taxon *Iainarchaeum andersonii* SCGC AAA011-E11: either at the Archaea root or as sister of the *Methanobacteriota* taxon *Thermococcus litoralis* DSM 5473. Therefore, we selected the untreated and the 20% site removal trees as representatives of these two topologies, respectively, and conducted molecular clock analysis in each case by applying the RelTime method, which has been popularly used for estimating divergence times under different evolutionary rates among lineages [59–62].

### Supplementary Methods

**Generation of the global non-redundant *Poseidoniales* genome dataset.**

To obtain potential brackish *Poseidoniales* genomes, we used IDBA-UD (v. 1.1.3) [63] to assembly clean reads of the metagenomes of the Pearl River estuary, Shenzhen Bay, the Brisbane River estuary, the Jiulong River estuary, the Yangtze River estuary, the Columbia River estuary, the Amazon River estuary, Caspian Sea, and Baltic Sea (table S1). Contigs longer than 2 kb were used for binning by using the binning module of MetaWRAP recruiting metaBAT2 [64], Maxbin2 [65], and CONCOCT [66] methods. Bins with completeness >50% and contamination <10% as evaluated by CheckM (v. 1.0.5) [67] were kept and those classified as *Poseidoniales* by GTDB-tk (v. 1.3.0, release 95) [68] were used for downstream analysis. Previously published marine *Poseidoniales* genomes generated by Rinke et al. [69], Tully [70], and Orellana et al. [16] were downloaded from their online deposits. The combination of downloaded genomes with those generated in this study results in 835 *Poseidoniales* metagenome-assembled genomes (MAGs) (table S2). Potential contaminant contigs in each MAG were further removed by manual check aided by acdc (v. 1.2.1) [71]. A non-redundant MAG dataset was generated by using dRep (v. 2.6.2) [72] and setting a cutoff of 99% average nucleotide identity. This dataset contains 455 *Poseidoniales* MAGs. Quality check and taxonomic classification of these MAGs were conducted by using CheckM and GTDB-tk, respectively. A manually selected subset containing 148 MGII-specific single copy marker genes (present in at least 328 of the 455 *Poseidoniales* MAGs) were selected for MAG completeness assessment when CheckM was used (table S5). Genes and proteins of the MAGs were predicted by using Prodigal (v. 2.6.3) [73].

**Phylogenomics of the non-redundant *Poseidoniales* MAGs.**

We used hmmsearch (v. 3.1b2; -E 1E-5) [74] to search for the 122 archaeal single-copy marker proteins [67] in the 455 non-redundant *Poseidoniales* MAGs based on hidden Markov models (HMMs) in Pfam [75] and TIGRfam [76] databases. MGIII euryarchaeal and other archaeal genomes were used as the outgroup (table S4). Marker proteins present in ≥60% taxa were retained and aligned, respectively, by using MUSCLE (v. 3.8.1551; --maxiters 16) [77]. The alignment matrixes were denoised by using trimAl (v1.2rev59; -automated1) [78] and then concatenated. Missing data were filled with gaps. A maximum-likelihood tree was reconstructed by using FastTree (v. 2.1.10; -gamma -lg) [79] and visualized in the Interactive Tree of Life (iTOL, v.5.1.1) [80].

**MAG abundance calculation.**

To profile the distribution of *Poseidoniales* in global marine surface water, the non-redundant MAGs were mapped by clean reads of metagenomes and metatranscriptomes obtained from surface samples of the Pearl River estuary, Shenzhen Bay, the Jiulong River estuary, the Yangtze River estuary, the Columbia River estuary, the Amazon River estuary, Caspian Sea, Baltic Sea, the Helgoland region, the Port Hacking offshore region, Northwest Pacific, and the Tara oceans project (table S1). To minimize potential unspecific mapping, rRNA and tRNA genes in the MAGs were identified by using Metaxa (v. 2.2) [81], and low complexity regions were predicted by using DustMasker (v. 1.0.0) (https://github.com/ncbi/ncbi-cxx-toolkit-conan). These regions of the MAGs were masked before mapping using Bedtools (v. 2.27.1). Read mapping was conducted by using Bowtie2 (v. 2.3.5) [82] and followed by sorting and format convert to BAM files by using SAMtools (v. 1.9) [83]. The BAM files were filtered by using BamM (v. 1.7.3) (https://github.com/minillinim/BamM;) with thresholds of 99% identity and 75% coverage.

Finally, bbmap (http://jgi.doe.gov/data-and-tools/bb-tools/) was used to calculate read counts for each contig and the RPKM (Reads Per Kbp of each genome per Mbp of each metagenomic sample) value was calculated for each MAG in each sample, respectively.

**Proteome acidity estimation.**

The isoelectric points (p*I*) of proteins of MAGs were calculated by using Pepstats of the EMBOSS package [84]. p*I* frequency distribution of a proteome was calculated as previously described [2]. Proteome acidity in this study is defined as the ratio of the frequency of the acidic peak (p*I* 4.5) to the frequency of the semi-acidic peak (p*I* 6.25) as shown in fig. S8.

**Habitat salinity range analysis.**

Habitat salinity was investigated by calculating the abundance (RPKM) of *Poseidoniales* MAGs in metagenomes from diverse salinities (table S1). A MAG is considered present in a metagenome if its RPKM value is above 0.01. The up-limit habitat salinity of a *Poseidoniales* taxon is set as the highest salinity where it is present, and the down-limit is set as the lowest salinity where it is present. Its optimum habitat salinity is set as the salinity where it has the highest RPKM value.

**Functional annotation and comparison of MAGs.**

Protein sequences of MAGs were annotated based on the KEGG database by using kofamscan[85], and the COG [86], arCOG [87], Pfam [88] and Tigrfam databases [76] by using BLASTp [89] (E-value < $10^{-3}$, bit score > 50, similarity > 50%, and coverage > 70%), respectively. Genes specifically enriched in brackish *Poseidoniales* were defined as those present in > 85% brackish MAGs but in < 5% marine MAGs. Genes specifically enriched in marine *Poseidoniales* were defined *vice versa*.

To compare proteome acidity values of *Poseidoniaceae* with and without *corA*, the F-Test was conducted in Microsoft® Excel for Mac (v. 16.69) and a P value of 0.002 was obtained suggesting the variances of the two groups are significantly different. The Student's *t*-Test was then conducted by setting a two-tailed distribution and type of two-sample unequal variance.

**Tree of *Poseidoniales* and other archaea.**

To build the tree for the dataset containing 188 taxa (fig. S7), 39 of the 41 marker proteins described by Adam *et al.*, 2017 [51] were used. The other two proteins were excluded because they are absent in most of the *Poseidoniales* MAGs in this dataset. Detection of the marker proteins in each MAG was based on functional annotation. The marker proteins of each MAG were identified according to the genome functional annotations. A multisequence alignment of concatenated marker proteins was constructed by using MUSCLE and automatically trimmed by using trimAl. Removal of compositional heterogenous sites was conducted by applying a $\chi^2$-score-based approach[90]. Maximum likelihood trees were reconstructed with the LG+C60+F model implemented in IQ-TREE (v. 2.0.3) [91] and then visualized in iTOL. Maximum-likelihood trees of the dataset containing 230 taxa (fig. S9) were reconstructed in the same manner.

**Amalgamated likelihood estimation (ALE) analysis.**

Functional genes in the 231-taxa dataset were aligned by using MAFFT L-INS-I [92] and denoised by using trimAl (automated1). The ML tree was constructed by using IQ-TREE with the parameters "-seqtype AA -m LG+PMSF+G -B 1000 --bnni". The ALEml_undated algorithm of the ALE package [93] was used to reconcile the functional gene tree against the phylogenomic tree to infer the numbers of duplication, loss, transfer (within the sampled genome set), and origination (including both transfer from other phyla outside the species tree or de novo gene formation) on each branch of the *Thermoplasmatota* species tree. The results were visualized in iTOL.

**Gene gain/loss event analysis for the BK4-BK5-M-BK6 monophyletic clade of *Poseidoniaceae*.**

The event number of a gene (KO or arCOG entry) in a terminal taxon (MAGs) is 1 if the gene is present and is 0 if absent. The event number of a gene in an internal node is defined as the DTLO event numbers calculated by applying the branchwise_numbers_of_events.py script described by Sheridan *et al.* [94]. Gene gain/loss events between adjacent internal nodes or between adjacent internal nodes and terminal taxa are defined as the following: 1) A loss event is defined if the event number of the older node (an internal node) is greater than 0.8 and is eight times greater than that of the younger node (an internal node or a terminal taxon); and 2) A gain event is defined if the event number of the younger node (an internal node or a terminal taxon) is greater than 0.8 and is eight times greater than that of the older node (an internal node).

**Molecular clock analysis and speciation rate calculation.**

Node divergence time of the 231-taxa maximum-likelihood trees was estimated by using RelTime in MEGA X (v10.1.5) with the LG+G model and with 95% confidence interval[59]. The root of Archaea (4.38-3.46 Ga) [95–97] and three constraints (i.e. the roots of *Thermoproteales*, *Sulfolobales* and *Thermoplasma*) related to the Great Oxygenation Event (2.32 Ga) [98,99] were used for calibration as introduced in our previous study[58]. The speciation rate of each branch in the timetrees were estimated by BAMM (v2.5.0) [100].

### Supplementary Figures

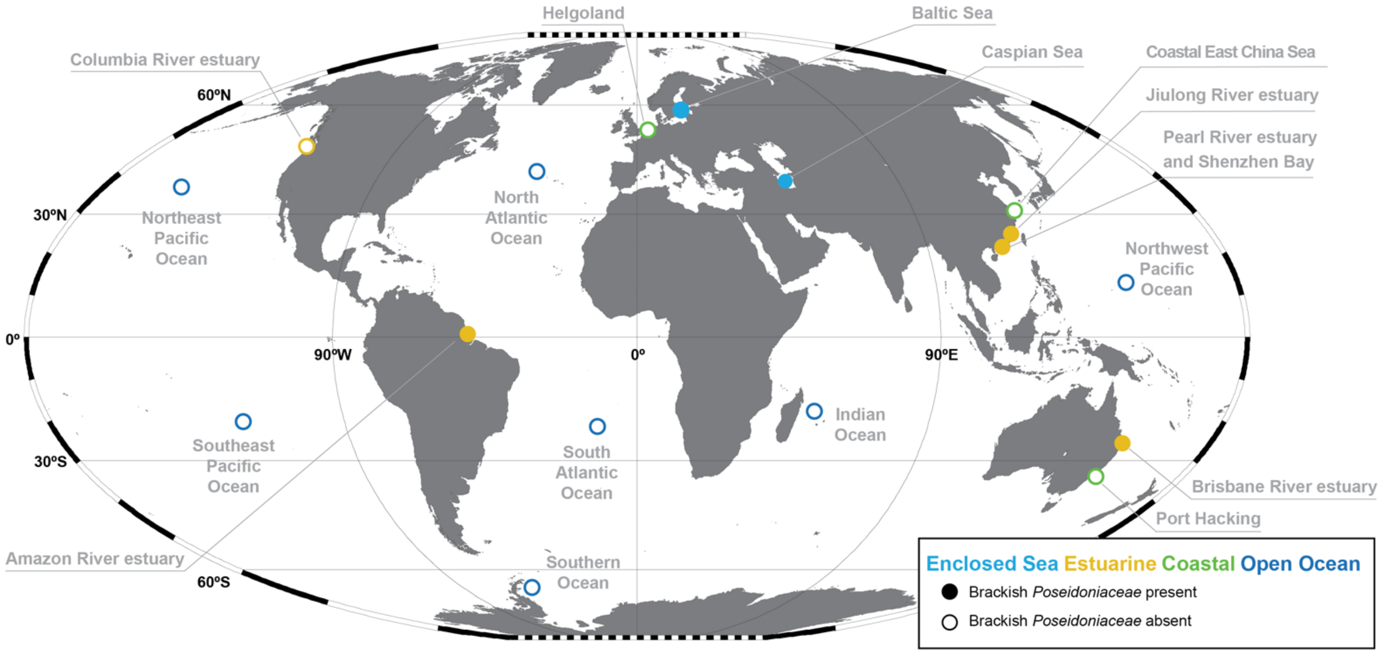


**fig. S1. Station map of the metagenome samples collected.**

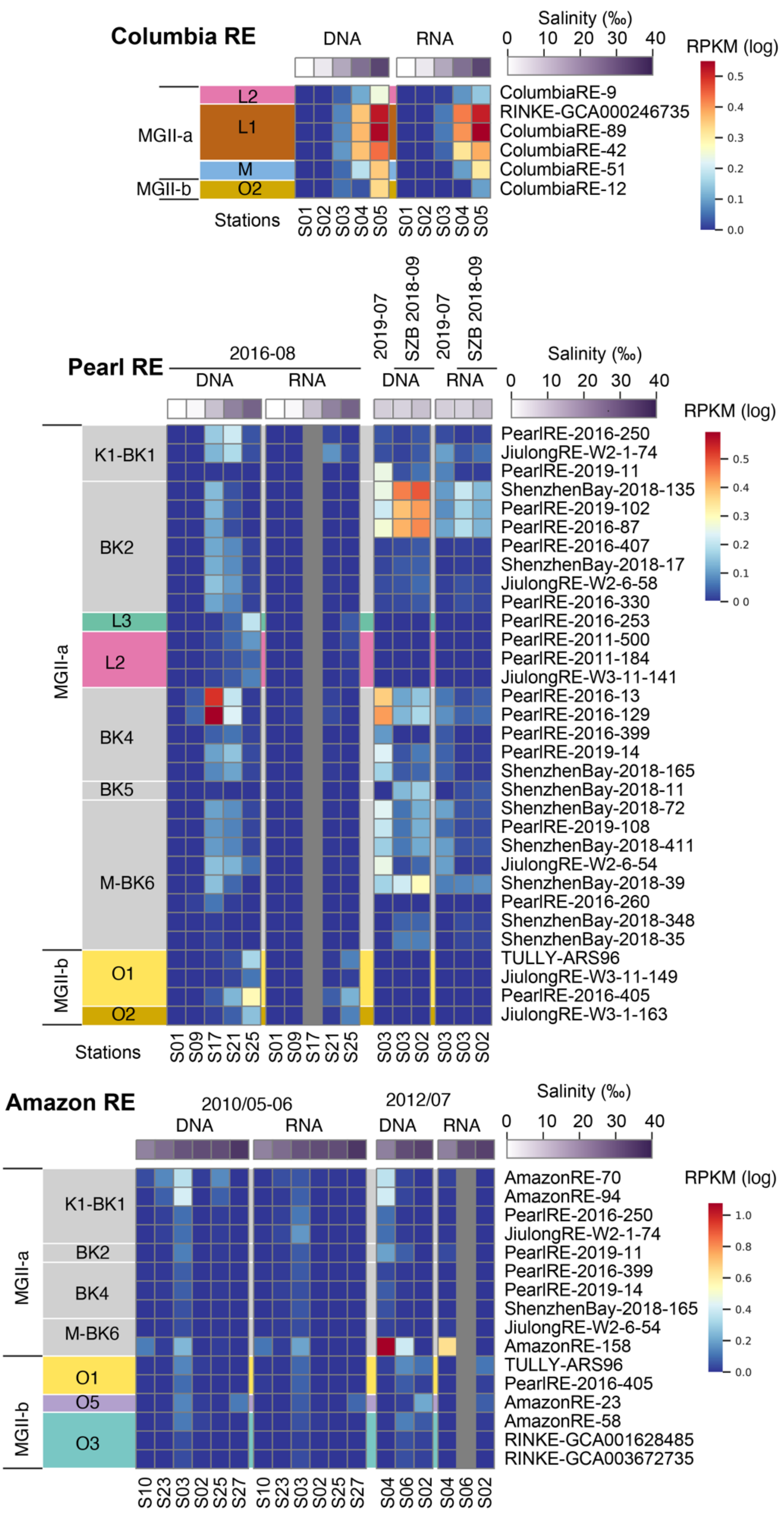


**fig. S2. Metatranscriptome read abundance of *Poseidoniales* MAGs at estuaries.** Abundance is calculated as reads per kilobase per millions of reads (RPKM). Shade on branches show *Poseidoniales* genera with the color code in consistent to Rinke et al. 2019 [69], except for brackish clades which are in grey.

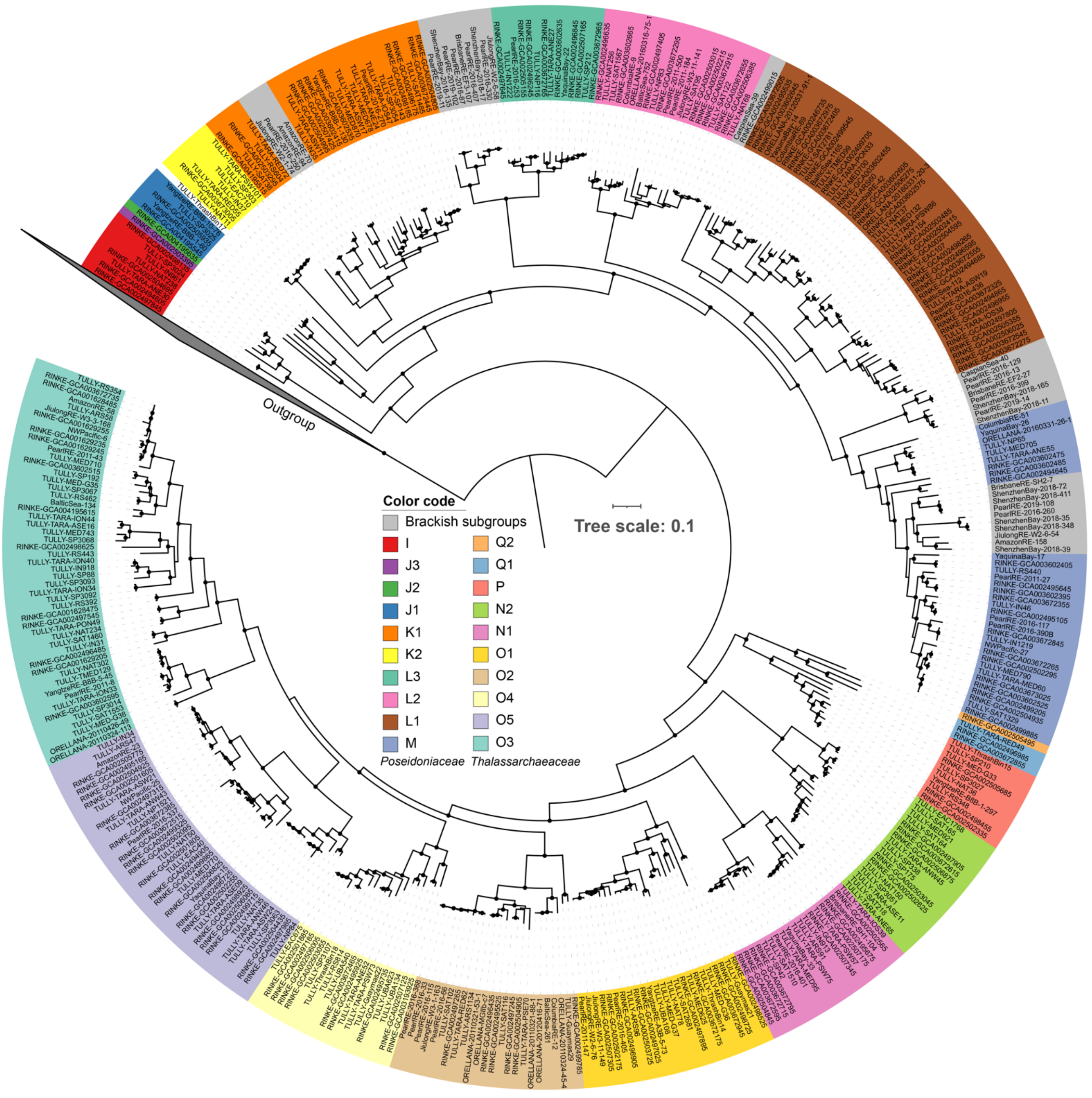


**fig. S3. Phylogenomic tree of genomes in the non-redundant *Poseidoniales* dataset.** Maximum-likelihood tree is shown. Shade on branches show *Poseidoniales* genera with the color code in consistent to Rinke et al. 2019 [69], except for brackish clades which are in grey. Bootstrap values (1000) >0.95 are shown as dots.

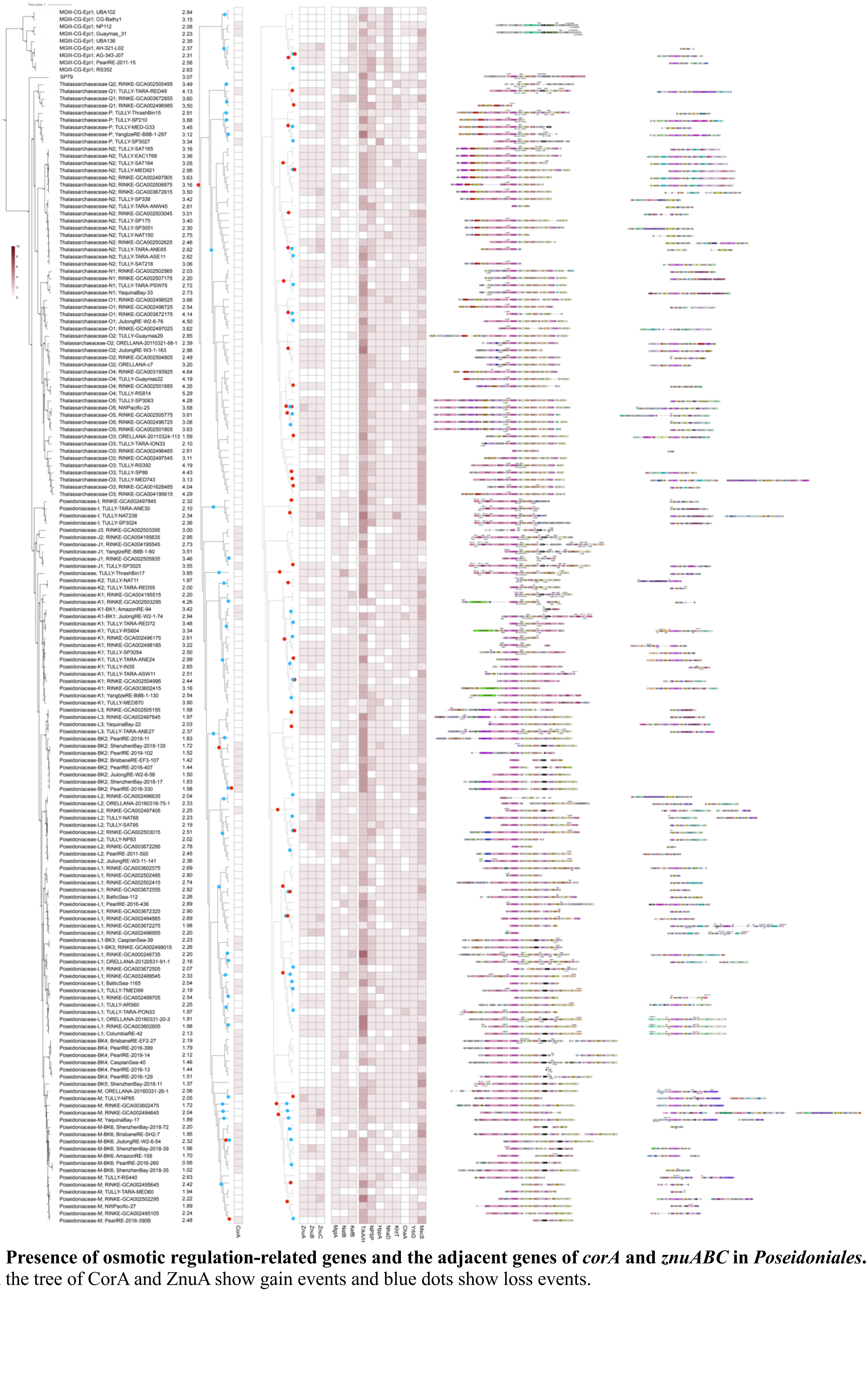

**fig. S4. Presence of osmotic regulation-related genes and the adjacent genes of *corA* and *znuABC* in *Poseidoniales*.** Red dots on the tree of CorA and ZnuA show gain events and blue dots show loss events.

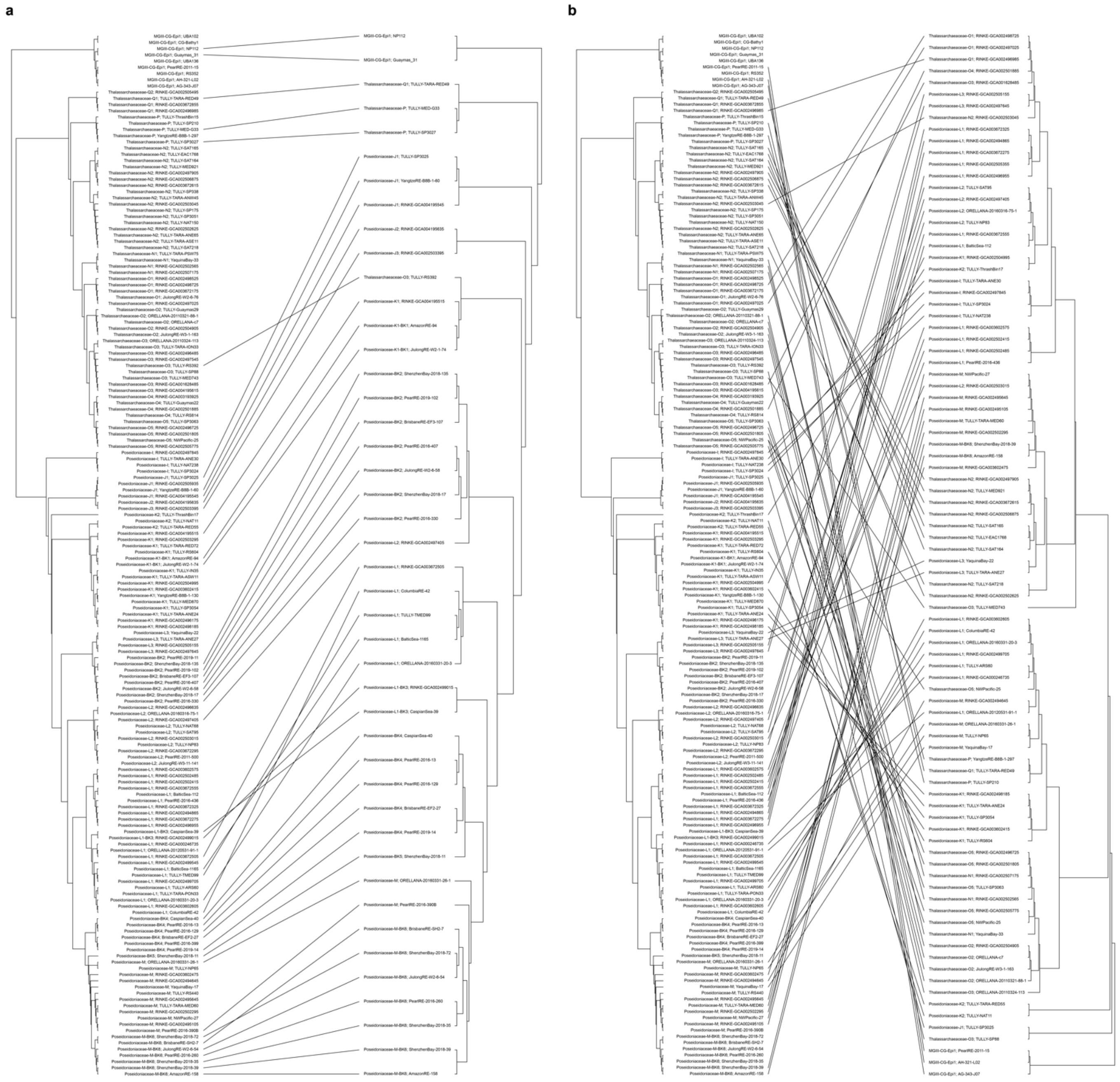


**fig. S5. Comparison of the genome tree and the trees of CorA (A) and ZnuA (B), respectively.** The genome tree is shown on the left of each panel. The gene trees of CorA and ZnuA are shown on the right of each panel, respectively. Lines connect the same taxa.

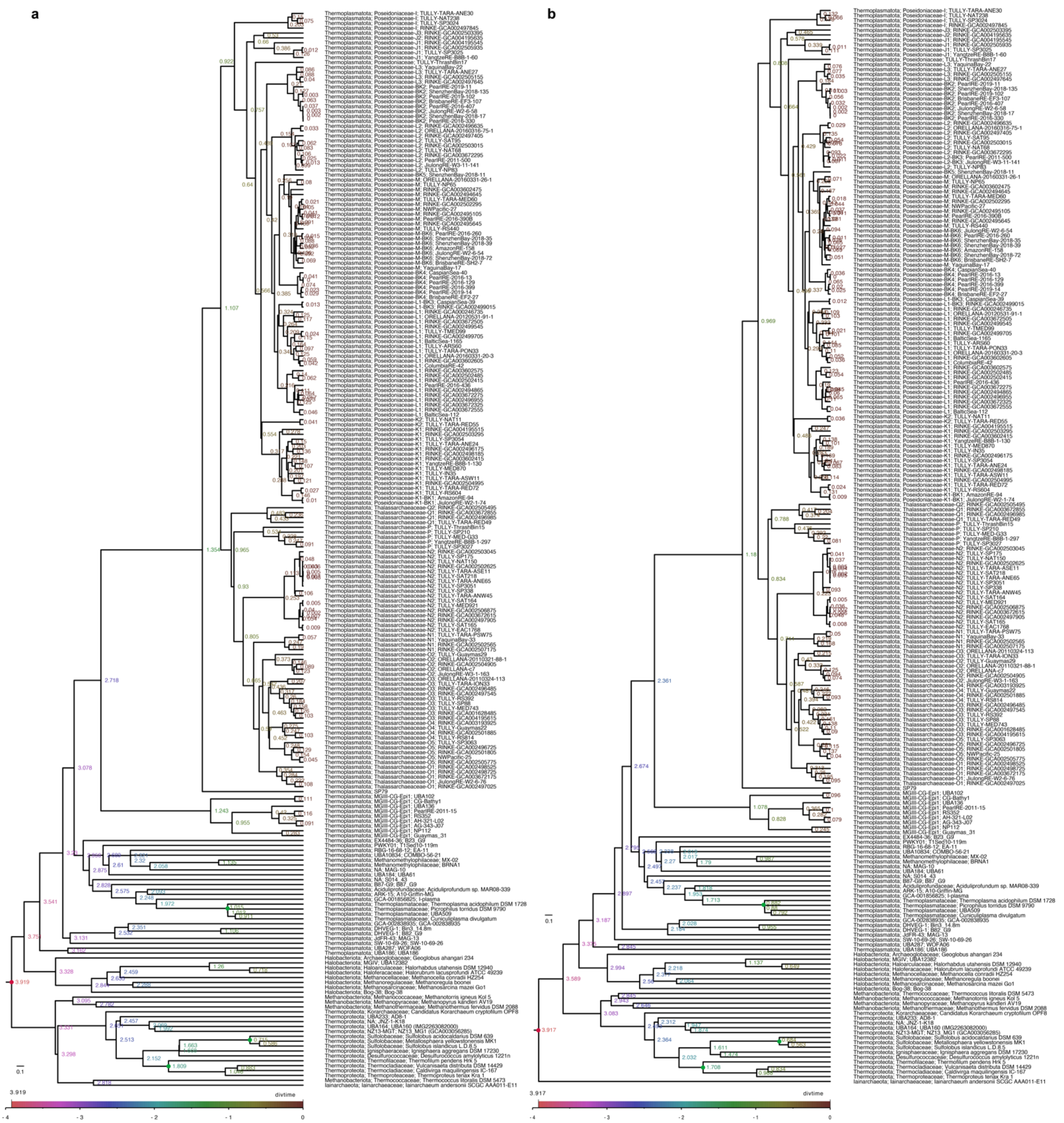

**fig. S6. Molecular dating of the phylogenomic trees of the 231-taxa dataset.** Trees of the untreated (**A**) and 20% most heterogeneous sites removed (**B**) are shown. Estimated divergence time is shown on nodes. The red dots show the root of Archaea and the green dots show three constraints related to the Great Oxygenation Event.

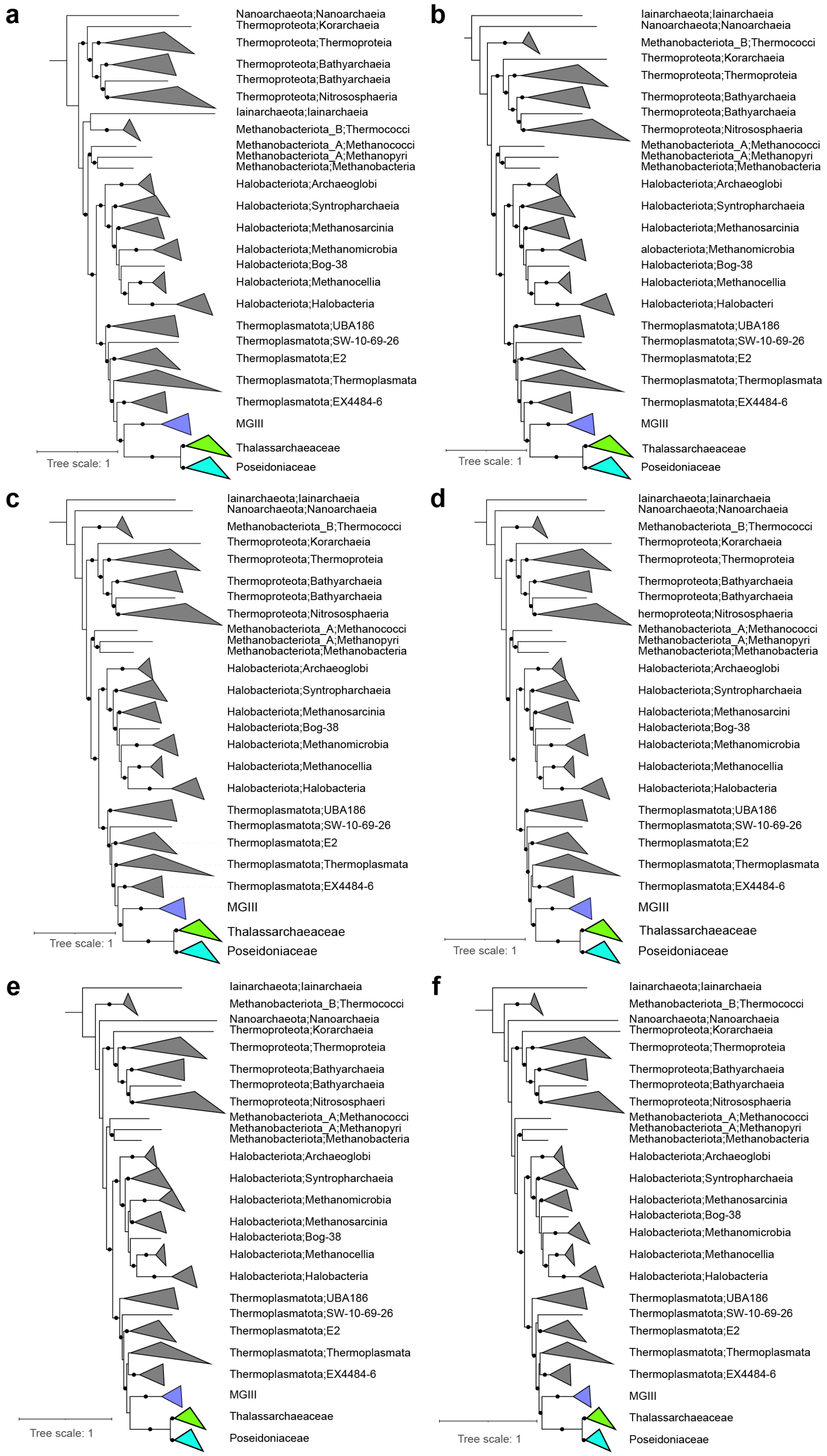


**fig. S7. Phylogenomic trees of the 188-taxa dataset.** In the maximum-likelihood trees of untreated (**A**), 5% (**B**), 10% (**C**), 20% (**D**), 40% (**E**) and 60% (**F**) most heterogeneous sites removed, solid dots on internal branches show branch supports of ultra-fast bootstrapping (1000) in IQTree > 90%.

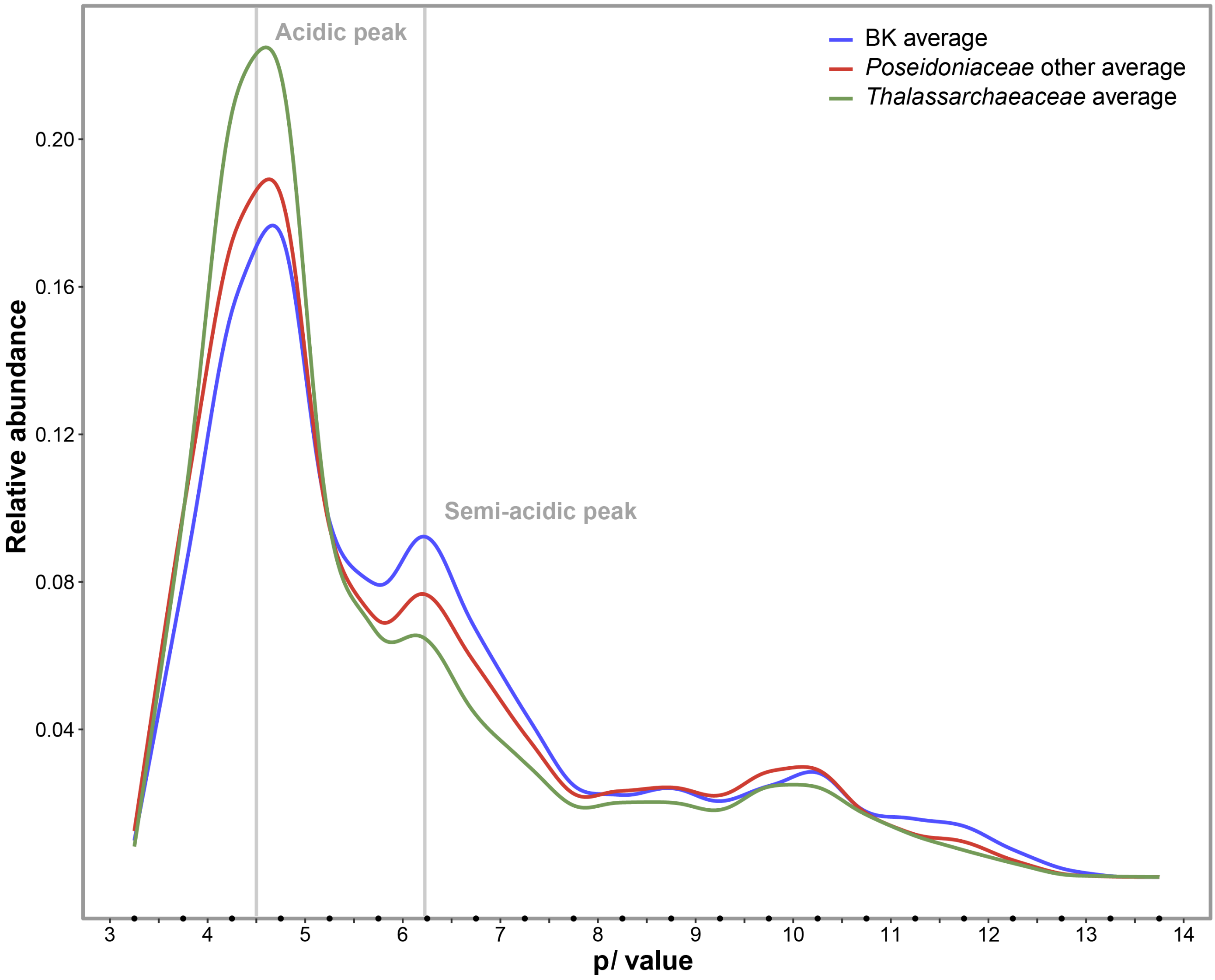


**fig. S8. p*I* distribution of brackish and marine *Poseidoniales* subgroups.**

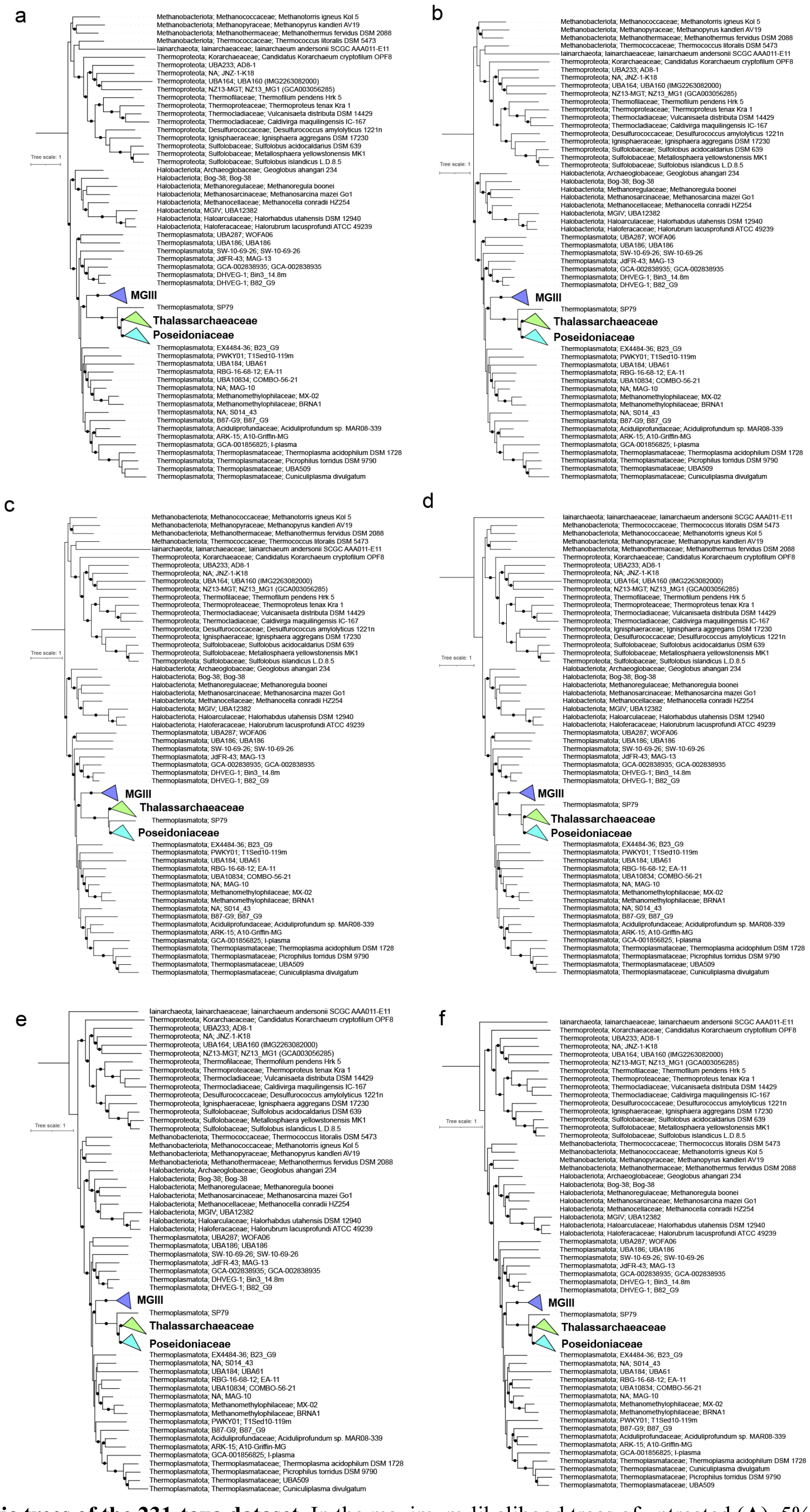


**fig. S9. Phylogenetic trees of the 231-taxa dataset.** In the maximum-likelihood trees of untreated (**A**), 5% (**B**), 10% (**C**), 20% (**D**), 40% (**E**) and 60% (**F**) most heterogeneous sites removed, solid dots show branch supports of ultra-fast bootstrapping (1000) in IQTree > 90%.
